# Outward-facing P-glycoprotein bound to drug displays nucleotide-dependent drug egress mechanism

**DOI:** 10.64898/2025.12.04.692479

**Authors:** Sungho Bosco Han, Hao Fan, Stephen M. Prince, Jim Warwicker

## Abstract

ABCB1/P-glycoprotein requires coordination between ligand movement in the trans-membrane cavity and ATP binding and hydrolysis at the nucleotide binding domains, yet the sequence of these events remains incompletely resolved. Here, atomistic simulations with paclitaxel and verapamil across outward facing P-glycoprotein ensembles in various nucleotide states were integrated with analysis of transmembrane domain contacts, geometric solvent/ligand pathways, extracellular gating distances, and nucleotide coordination characterization. Collectively, our simulations delineate a mechanistic framework for P-glycoprotein in which ATP binding organizes an outward-facing ensemble and sequential hydrolysis at the two nucleotide-binding sites drives substrate progression toward egress. Asymmetry in the nucleotide state is correlated to distinct motions within the transmembrane domains, which suggests a possible transient extracellular gating mechanism that enables staged release and limits re-entry of the ligand. The resulting model connects nucleotide chemistry to transmembrane domain motions that regulate gate dynamics and may guide experiments and development of modulators for ABC exporter activity in physiological and pharmacological contexts.

## 1 Introduction

The human P-glycoprotein (P-gp, ABCB1/MDR1) is a 170-kDa ATP-binding cassette (ABC) efflux transporter that plays a central role in exporting diverse xenobiotics and drugs from cells [1, 2]. P-gp is widely expressed in barrier tissues (intestinal epithelium, liver, kidney, brain endothelium, etc.) where it lowers intracellular drug concentrations, and its broad substrate spectrum (drugs from 100–4000 Da) underlies multidrug resistance in cancer and other diseases [3]. Structurally, P-gp consists of two homologous halves, each containing a transmembrane domain (TMD) and a nucleotide-binding domain (NBD), which together undergo large conformational changes powered by ATP binding and hydrolysis. Inward-facing (IF) structures of ligand-bound P-gp have been known for many years, but only recently has an ATP-bound, outward-facing (OF) conformation been determined by cryo-EM [4]. In that 3.4 Å structure of human P-gp, the NBDs form a tight dimer with two ATP molecules sandwiched between them, and the drug-binding cavity is re-oriented toward and open to the extracellular space, while being not accessible to the cytoplasm. Likewise, the first crystal structure of the bacterial homologue Sav1866 captured an outward-facing conformation (ATP-bound NBDs closed) with a central cavity exposed to the extracellular solvent [5].

Despite the increasing number of structural studies of P-gp and ABC transporters being conducted, the exact coupling mechanism between substrate translocation and the ATP hydrolysis cycle in P-gp remains incompletely understood. In particular, the detailed timing and sequence of substrate incorporation into binding cavity, nucleotide binding, ATP hydrolysis and unbinding of hydrolyzed products and substrate release still remains a mystery. Recent biochemical and biophysical studies have attempted to shed some light in these mechanistic details of the substrate translocation cycle, by capturing transient intermediate states in various methods. Electron paramagnetic resonance (DEER) spectroscopy on human P-gp showed that trapping the transporter in a post-hydrolysis transition-state (ADP-vanadate) stabilizes a short-lived outward-facing conformer, suggesting that ATP hydrolysis and phosphate release may drive the power stroke to expel substrate [6]. In this work, P-gp shows patterns that support a two-stroke “alternating sites” cycle, with observation that there is significant asymmetry between the two NBDs in the OF P-gp bound to drugs [7]. By contrast, the cryo-EM structure of ATP-bound P-gp E-to-Q mutant (catalytically inactive glutamate mutants) showed an outward-facing state with no substrate present, supporting a model where substrate extrusion can precede hydrolysis [4]. Accordingly, substrates and inhibitors have been shown to differentially affect the conformational cycle of P-gp: substrates accelerate ATP turnover by stabilizing an asymmetric post-hydrolysis state, whereas high-affinity inhibitors favor a symmetric, substrate-free cycle [7]. Other ABC transporters suggest a “twist-and-squeeze” mechanism, in which ATP binding (even without hydrolysis) induces a global twisting contraction that expels the bound compound through transient gaps in the TMDs [8]. These studies underline open questions such as when exactly the substrate leaves the TMD cavity, whether one or two ATP molecules must be hydrolyzed for transport, and how P-gp ensures that the drug does not back-diffuse before the channel resets.

Another key area that remains unclear is how the ligand egress pathways in P-gp are formed and controlled. High-resolution cryo-EM structures of human P-gp with substrate bound (e.g. vinblastine, verapamil) revealed multiple IF conformations with varying separation distance between NBD dimer where the cavity access notably differed, showing that P-gp can adapt multiple conformations in IF state that may affect the drug packing within the cavity [9]. Another cryo-EM study utilized mouse P-gp bound to AAC through a disulfide bond, which revealed not only IF and OF structures but multiple occluded intermediate structures that had the translocation cavity closed at both ends [10]. This occluded state sits between the substrate-bound inward form and the post-release closed form, suggesting it accommodates the drug as the NBDs dimerize. In another study, narrow tunnels around TM1, 6, 7, and 12 were identified, implying that drug exit may occur through limited “cracks” rather than a large open vestibule [8]. Nasim *et al.* showed that a pair of symmetric tyrosine residues (Y310 and Y953) at the extracellular apex act as two gates controlling release of specific substrates [11]. Other work has mapped clusters of aromatic and hydrophobic residues lining the central cavity (e.g. F303, F335, Phe978) that make frequent contacts with diverse substrates [8]. It remains unclear, however, whether substrates versus inhibitors exit via the same route. Multiple cryo-EM structures revealed that small molecule inhibitors (e.g. elacridar, zosuquidar) can occupy overlapping sites with substrates but often stabilize distinct transporter conformations in IF state [12, 13]. A key goal is thus to delineate the molecular triggers (e.g. specific NBD or TMD residues) that couple nucleotide binding/hydrolysis to opening of the extracellular gate, and to determine how occupancy of the binding cavity by different ligands biases these events.

The outward-facing conformations of P-gp can be classified into several sub-states. In the classic outward-open state (as seen in the bacterial Sav1866 and human ATP-bound P-gp), the NBDs are fully dimerized around two ATP molecules, and the TM cavity is inaccessible from the intracellular side but open to the extracellular milieu [5]. By contrast, occluded states (observed by cryo-EM and spectroscopic methods) are thought to have both the cytoplasmic and extracellular gates closed around the substrate [10]. For example, Gewering *et al.* captured an occluded conformation that appears prior to ligand release, as the disulfide-linked AAC molecule bound inside the TMD cavity with occluded outer gate [10]. And lastly, we expect the protein to adapt a closed conformation once the drug is released to the extracellular space to avoid back-transportation of any molecule while preparing for the conformational reset back to the IF state. Kim *et al.* showed an ATP-bound P-gp structure in this OF closed conformation, in which the drug-binding pocket is oriented toward the extracellular solvent but constricted to preclude ligand binding [4]. These intermediate conformers likely represent steps in the transport cycle: an OF-occluded state accommodating the substrate as the NBDs close, then a possible OF-open state allowing release of bound drug, followed by a fully collapsed OF-closed state with the ligand still in transport. Structurally describing how the TMD helices rearrange between these OF-occluded, OF-open, and OF-closed states is critical for understanding how a substrate can move from the inner cavity to the extracellular space. In this work, we investigate how nucleotide binding at the NBDs and drug occupancy in the TMD jointly shape the outward-facing conformational landscape of human P-gp. Building on our prior drug-free simulations of IF P-gp states (Chapter 2), we use all-atom molecular dynamics of realistic human P-gp models embedded in a lipid bilayer. We examine multiple nucleotide conditions together with two different ligands, paclitaxel and verapamil docked within the TMD cavity. By comparing these systems in three different initial outward conformations (OF-occluded, OF-open, OF-closed), we aim to determine how NBD state and ligand identity co-modulate the TMD geometry and ligand dynamics.

## 2 Results

Full-length models of human P-glycoprotein in OF conformation were constructed and embedded in a POPC/cholesterol membrane environment as the basis for atomistic simulations. Using a standardized protocol [14, 15], we performed all-atom molecular dynamics to characterize the conformational behavior of P-gp in the presence of drug molecule (paclitaxel or verapamil) in four different nucleotide conditions, ADP/ADP, ADP/ATP, ATP/ADP and ATP/ATP; full methodological details are provided in Materials and methods.

**Table 1:** Total number and length of initial 10 ns multi-replica MD simulations performed with drug-bound OF P-gp. All replicas were initialized from the same starting configuration with a random seed of initial velocities.

| Drug | Paclitaxel |  |  | Verapamil |  |  |
| --- | --- | --- | --- | --- | --- | --- |
| Conformation | occluded | open | closed | occluded | open | closed |
| No. of nucleotide system | 4 | 4 | 4 | 4 | 4 | 4 |
| No. of replica simulation | 120 | 120 | 120 | 120 | 120 | 120 |
| Run time (ns) | 10 | 10 | 10 | 10 | 10 | 10 |
| Total run time (μs) | 4.8 | 4.8 | 4.8 | 4.8 | 4.8 | 4.8 |

**Table 2:** Total number and length of extended multi-replica MD simulations performed with drug-bound OF P-gp. Replicas showing outlier structural deviations in any predefined P-gp subdomain (see Methods) were extended to 100 ns from the initial 10 ns production run.

| Drug | Paclitaxel |  |  | Verapamil |  |  |
| --- | --- | --- | --- | --- | --- | --- |
| Conformation | occluded | open | closed | occluded | open | closed |
| No. of nucleotide system | 4 | 4 | 4 | 4 | 4 | 4 |
| No. of replica simulation | 25 | 28 | 19 | 26 | 29 | 21 |
| Run time (ns) | 100 | 100 | 100 | 100 | 100 | 100 |
| Total run time (μs) | 10 | 11.2 | 7.6 | 10.4 | 11.6 | 8.4 |

### 2.1 Ligand contact profile in various nucleotide conditioned P-gp in occluded conformation

In the occluded state, the drug-binding cavity is enclosed toward the extracellular side, based on the mouse P-gp structure with disulfide-linked AAC peptide, showing the ligand residing in mid-cavity with no solvent exposure [10]. We docked one of the two drug molecules in the same binding pocket occupied by AAC peptide ligand, and both paclitaxel and verapamil contacted residues deep in the chamber (Fig. 2g,h). Notably, the pre-hydrolytic (ATP/ATP) condition yielded the fewest high-frequency contacts for both drugs, suggesting a looser initial binding. This is evident for paclitaxel, which in ATP/ATP condition, engaged only 1̃1 residues with >25% contact probability (e.g. Q195 in TM3 and F343 in TM6; Fig. 2a), compared to 1̃6–17 contacts in post-hydrolytic states (ATP/ADP, ADP/ATP, ADP/ADP; Fig. 2c,e,g). Verapamil showed an analogous trend: only 11 residues were involved in ATP/ATP (Fig. 2b), whereas ATP/ADP and other ADP-bound states engaged 1̃5–16 contacts (Fig. 2d,h). Overall, our data indicates nucleotide states representing hydrolyzed conditions correlate with a broader drug–cavity interface in the occluded conformation (Fig. 1a,b).

**Figure 1:**
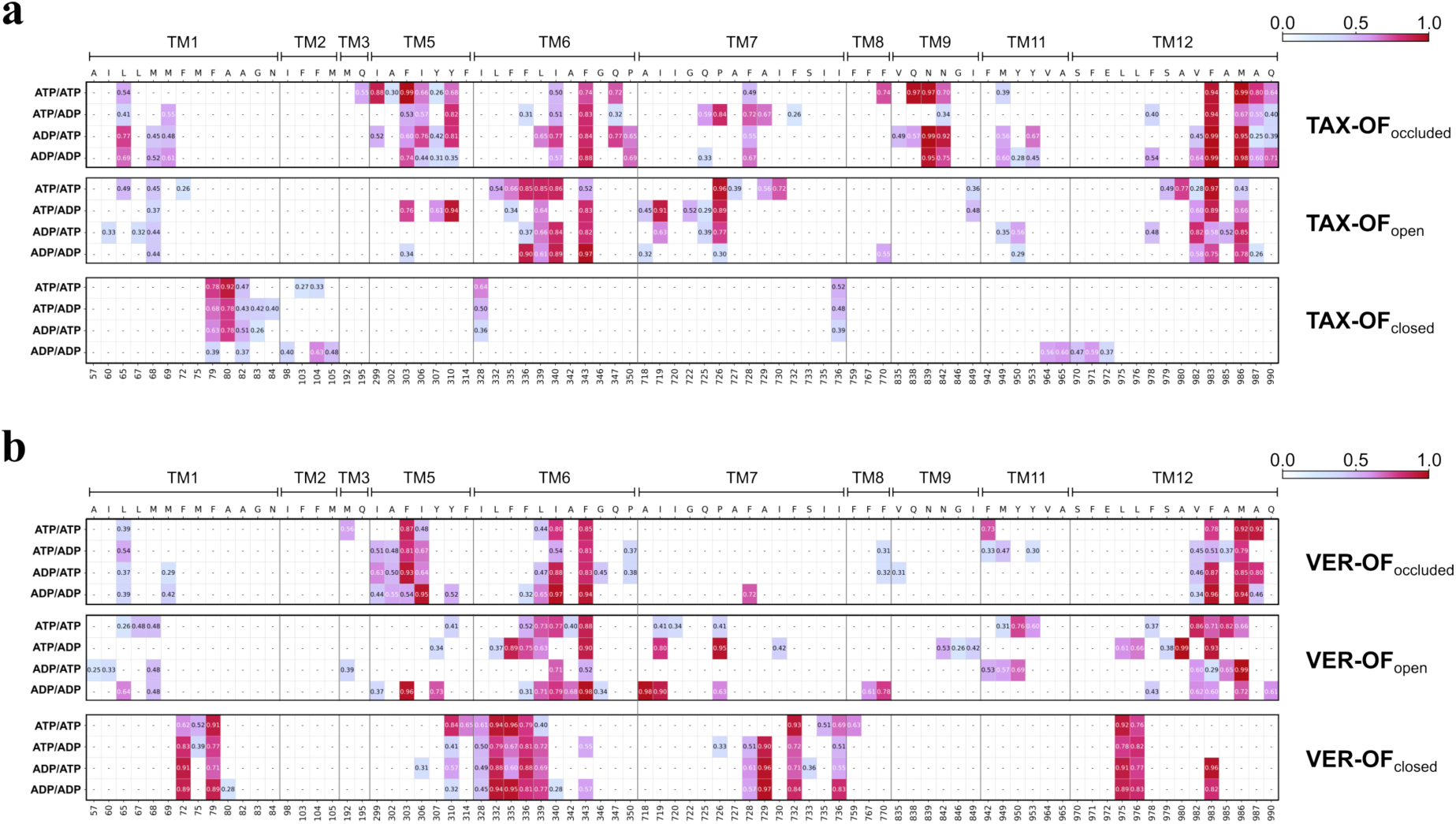
Paclitaxel/Verapamil-TMD interaction profile in OF P-gp. Either paclitaxel or verapamil was docked in OF-occluded (human homology model built based on mouse L335C mutant OF occluded cryo-EM structure), OF-open (human homology model built based on SAV1988 OF X-ray crystal structure 2HYD [5]), and OF-closed P-gp (human homology model built based on human OF E-to-Q mutant cryo-EM structure 6C0V [4]). (a) The interaction frequency (with paclitaxel) of the protein residues are shown as heatmap for each of the three OF P-gp systems initially docked with paclitaxel in OF-occluded, OF-open and OF-closed conformations. For each of the three systems, the interaction frequency heatmaps are shown for each of the four nucleotide states (ATP/ATP, ATP/ADP, ADP/ATP and ADP/ADP) P-gp systems in each row. (b) The interaction frequency (with verapamil) of the protein residues are shown as heatmap for each of the three OF P-gp systems initially docked with verapamil in OF-occluded, OF-open and OF-closed conformations. For each of the three systems, the interaction frequency heatmaps are shown for each of the each for four nucleotide states (ATP/ATP, ATP/ADP, ADP/ATP and ADP/ADP) P-gp systems in each row.

**Figure 2:**
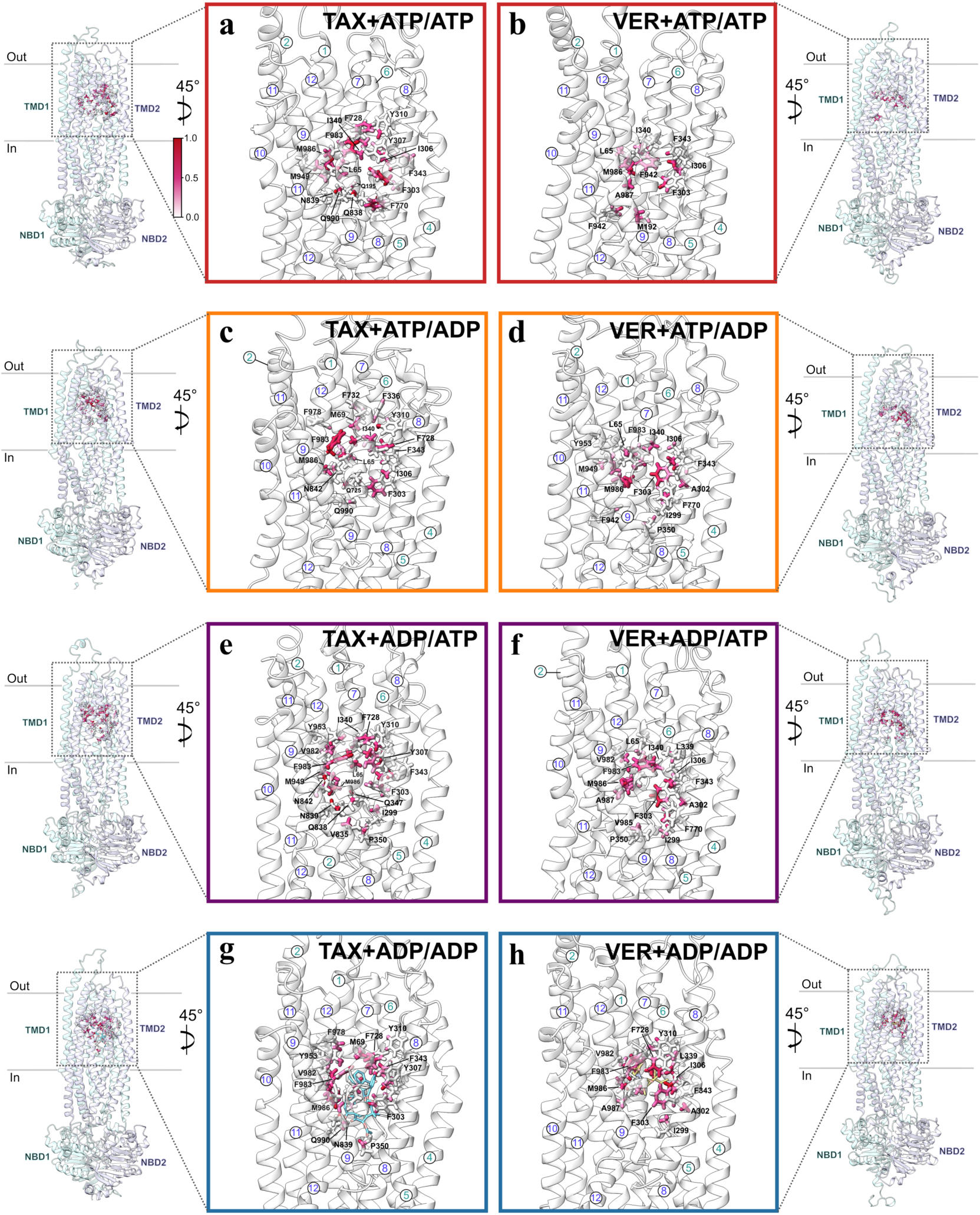
Residue level interaction frequencies between the TMD cavity residues and bound drug, either paclitaxel (TAX) or verapamil (VER) in OF-occluded P-gp. (a-h) On the side, it shows a global orientation/representation of OF-occluded P-gp to the side, with a box showing the clusters or TMD residues involved in the interaction with respective drug. The area shown with the box is displayed in an enlarged picture, where the residues with at least one atom above interaction frequency of 0.25 are shown with the side chain/backbone atoms colored in the respective frequency value. The initial docking position of TAX is shown in blue, in stick representation. Paclitaxel was docked slightly further inside the TMD ligand binding cavity near F303, F343, F983 side chains without access to extracellular solvent. (h) P-gp bound to VER with ADP/ADP. The initial docking position of VER is shown in yellow, in stick representation. Verapamil was docked slightly further inside the TMD ligand binding cavity near F303, F343, F983 side chains without access to extracellular solvent.

#### Paclitaxel in OF-occluded conformation

Paclitaxel consistently interacted with a broad swathe of the cavity, contacting residues from both halves of P-gp. Key contacts occurred on TM5, 6, 7, 9 and 12. For instance, F343 (TM6) and Q986 (TM12) were among the highest-frequency contacts in every state, underscoring the central role of these helices in paclitaxel binding (Fig. 2a, c–g). Despite this consistent contact, subtle differences emerged with nucleotide state. In the ATP/ATP condition (Fig. 2a), paclitaxel lacked certain TM1 contacts (M68, M69) that became prominent in ADP-bound states. Instead, only in ATP/ATP state, unique contacts were observed with Q195 on TM3, I299 and A302 on TM5, and F770 at TM8, which were all located at lower parts of the cavity. The additional involvement of TM3 and 8 suggests that prior to hydrolysis the drug cavity tightens from both sides of the cavity from the bottom (opposite of the extracellular space) possibly extruding drug toward the extracellular direction. In ADP consisting states representing the partially or fully hydrolyzed conditions, the drug shifted slightly: both asymmetric nucleotide conditions (ADP/ATP and ATP/ADP) lost many contacts near the bottom of the cavity seen in ATP/ATP state. In ATP/ADP condition in particular (representing NBS2 in post-hydrolytic condition), paclitaxel distributed further in the extracellular direction as it formed new contacts including F336 30% (TM6), F732 26% (TM7) and F978 40% (TM12). P-gp in ADP/ATP (NBS1, post-hydrolytic) state showed additional contacts with paclitaxel in all sides of the cavity, including additional residues closer to extracellular side (Y953 67%, V982 45%) and in bottom of the cavity (P350 65%, V835 39%). These shifts indicate that the position of paclitaxel adjusted with the nucleotide asymmetry, although the overall number of contacts remained comparable (1̃6 in both ATP/ADP and ADP/ATP). The ADP/ADP state (representing fully hydrolyzed condition) led to paclitaxel contacts in a distributed manner, involving a mixed set of residues seen in the asymmetric nucleotide conditions (engaging TM1, 5,6 while maintaining TM9, 11, and 12 contact). Overall, these data suggests that in the occluded conformation paclitaxel was deeply embedded and made extensive cavity contacts in ATP/ATP condition, and ATP/ADP condition notably favors drug to locate further toward extracellular side of the cavity. In other nucleotides, paclitaxel establishes slightly different set of residues to either distribute toward or away from the extracellular exit point without a clear directionality. The multi-ring structure of paclitaxel appeared to facilitate this adaptability (Fig. S4.2) : the aromatic rings maintained strong *π*–*π* or hydrophobic interactions with the cavity (notably against TM6 and TM12), while polar moieties (such as ester and hydroxyl groups) engaged surrounding residues without a strict nucleotide-state preference. For example, the phenyl rings of paclitaxel showed high contact frequencies with TM6 (L339/F343) and TM12 (A987) side chains, whereas the acetyl and benzoyl oxygen (ester carbonyls) and the taxane core (oxetane ring) formed intermittent contacts that persisted across states.

#### Verapamil in OF-occluded conformation

Verapamil exhibited a broadly similar interaction profile in the occluded cavity, albeit with fewer contact points consistent with smaller size. In ATP/ATP state, verapamil was nestled mid-cavity with contacts to only 10–11 residues, primarily on TM5, 6, 11 and 12 (Fig. 2b). In the ADP containing conditions, the ligand made more numerous and widespread contacts (1̃5 or more residues in each ADP-containing state). Like paclitaxel, verapamil engaged a core set of residues regardless of nucleotide state: I340 and F343 (TM6) and Q983/986 (TM12) remained frequent contact points (Fig. 1b). Interestingly, when ADP was incorporated into either NBS, the drug also consistently interacted with the TM5 region closer to intracellular side (I299, I302 44-63%) and additional residues in the mid-cavity (V982 3̃4-46%), reflecting broader contact distribution in the central pocket. In ATP/ADP condition in particular (Fig. 2d) showed verapamil contacting additional TM11 residues (M949 0.47%, Y953 0.30%) as the drug shifts slightly to the side in the mid-cavity. Conversely, in ADP/ADP (Fig. 2g), verapamil notably contacted residues further in the direction of extracellular side as it formed additional contacts with Y310 (52%) and F728 (72%), suggesting the drug distributes further down the extracellular exit route when both occupied nucleotides are in hydrolyzed forms. Ligand–moiety analysis indicated that the two aromatic rings of verapamil drove most of the hydrophobic contacts, sandwiching between TM6 and TM11/12, while the tertiary amine and nitrile group stayed near each side of the cavity (Fig. S4.1). The methyl ends of the dimethoxy moieties made persistent contacts with hydrophobic residues on the side of the cavity (TM1 and 5), but these did not significantly differ between nucleotide states. This suggested that verapamil, like paclitaxel, bound in a largely pre-formed pocket that was occupied in the covalently linked AAC ligand in the experimental structure [10], with modest readjustments in the presence of ADP that slightly broadened the interaction footprint.

### 2.2 Ligand contact profile in open conformation

The open outward-facing conformation, modeled based on the Sav1866 exporter structure, features a wide open cavity accessible from the extracellular space [5]. In the initial model structure, the drug-binding cavity is open to the solvent toward the extracellular space, and docked ligands sat near the upper chamber, partially exposed to solvent interface (Fig. 3g,h). In this state, TM helices of P-gp formed an expanded cradle around the ligand. Overall, both paclitaxel and verapamil maintained strong interactions with the cavity, particularly TM6 and TM12, while also contacting elements of the extracellular opening. The pre-hydrolysis ATP/ATP condition supported extensive drug contacts, whereas incorporation of ADP induced a shift in ligand positioning. Unlike OF-occluded system, both drugs bound in OF-open P-gp displayed a pronounced nucleotide asymmetry effect in the open conformation, wherein the ligand shifted toward the lobe containing the ADP-bound NBD, altering the extent of contacts.

**Figure 3:**
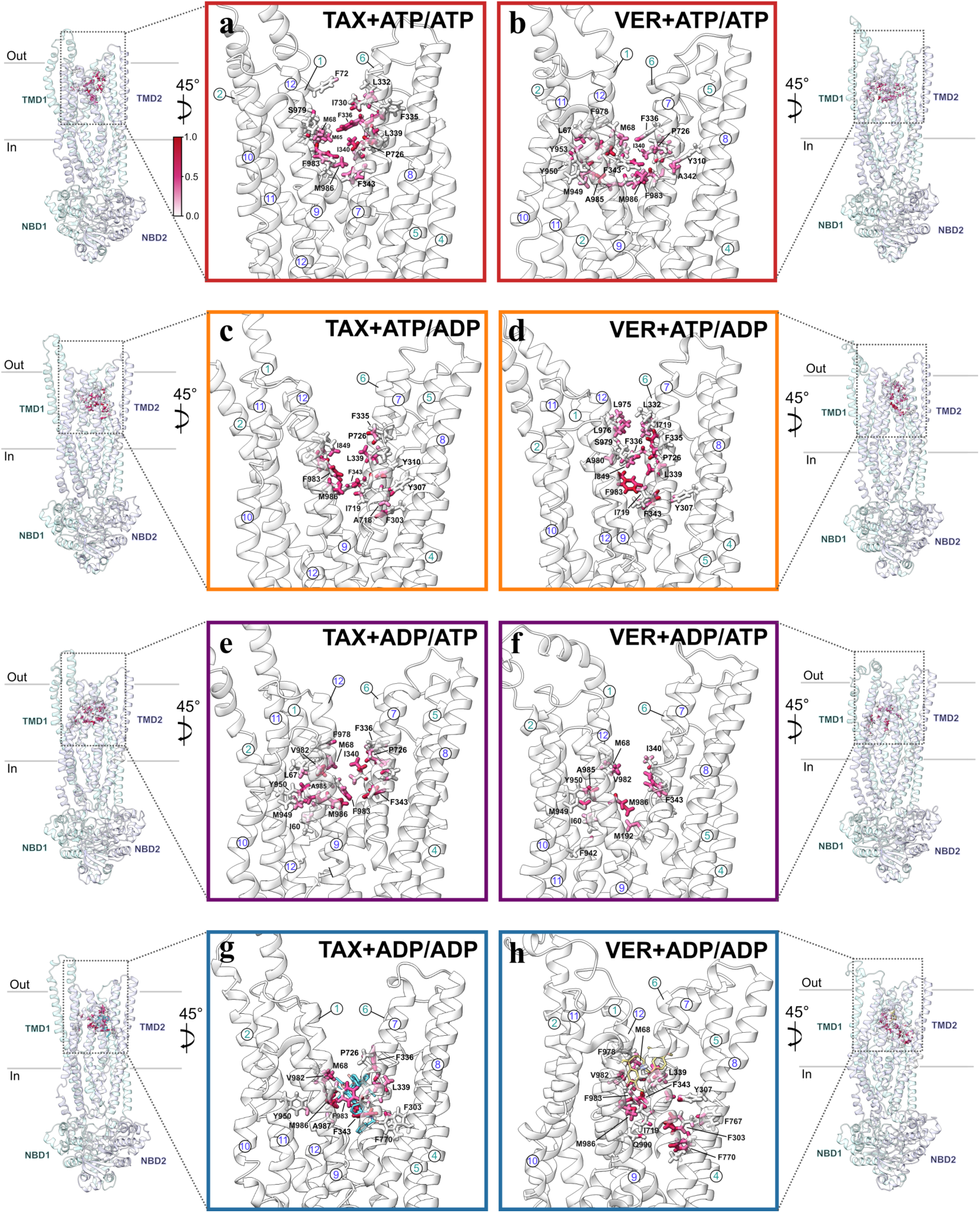
Residue level interaction frequencies between the TMD cavity residues and bound drug, either paclitaxel (TAX) or verapamil (VER) in OF-open P-gp. Panels a-h each shows a global orientation/representation of OF-open P-gp to the side, with a box showing the clusters or TMD residues involved in the interaction with respective drug. The area shown with the box is displayed in an enlarged picture, where the residues with at least one atom above interaction frequency of 0.25 are shown with the side chain/backbone atoms colored in the respective frequency value. The initial docking position of TAX is shown in blue, in stick representation. Paclitaxel was docked inside/top portion of the TMD ligand cavity near F343 and F983 side chains. (h) P-gp bound to VER with ADP/ADP. The initial docking position of VER is shown in yellow, in stick representation. Verapamil was docked inside/top portion of the TMD ligand binding cavity near F343 and F983 side chains.

#### Paclitaxel in OF-open conformation

In the open conformation, paclitaxel was positioned closer the extracellular end of the cavity. Consistent with this, paclitaxel in ATP/ATP made contacts that spanned from the top of TM1, 6 and 7 down to the midpoints of TM6 and TM12 (Fig. 3a). For example, in the ATP/ATP state (Fig. 3a). paclitaxel touched F72, L332/F335/F336 and I730 on the extracellular side of TM1, 6 and 7, which was unique to this pre-hydrolytic condition. Interestingly, the paclitaxel bound OF-open did not show any significant shift in drug contact distribution in the vertical direction (toward or away from the extracellular exit). Instead, the asymmetric post-hydrolysis conditions led to paclitaxel locating in the cavity on the side of NBS bound to ADP (Fig. 3). As the side picture of the entire P-gp shown in Figure 4.3, paclitaxel in OF-open P-gp with ATP/ADP and ADP/ATP shifted markedly to the side of cavity, showing that the cavity in OF-open conformation is sensitive to the chemical difference between ATP and ADP. ATP/ADP state (Fig. 3c) resulted in paclitaxel contacting additional TM5 and 7 residues as the drug shifts toward TM5/7 side of the cavity which is located on the side of NBD2 that is bound to ADP. Similarly, the ADP/ATP condition (Fig. 3e) led paclitaxel to interact with additional residues on TM11 as the drug shifts toward TM11 side of the cavity located on the same side as NBD1 which is ADP bound. In ADP/ADP condition (Fig. 3g), these additional contacts on TM5, 7 and 11 are all observed, as paclitaxel remains stably bound in the cavity interacting with all sides of the cavity. Throughout all open-state simulations, the R1 aromatic ring and taxane core of the drug continued to contact hydrophobic residues within the cavity while flexible substituents of paclitaxel (acetate ester and benzamide moieties) formed both hydrophobic and electrostatic interactions with the cavity (Fig. S4.2a). Despite shifts across nucleotide states, the overall binding orientation of paclitaxel remained such that the bulky bivalent shape of the drug was accommodated by the lower half of the cavity, with partial exposure at the portal in the R2 and R3 aromatic rings of the drug.

#### Verapamil in OF-open conformation

Binding of verapamil in the open conformation was likewise centered near the extracellular opening, but with even more pronounced sensitivity to nucleotide asymmetry. In the ATP/ATP state, verapamil made a broad array of contacts in the mid-cavity, totaling 1̃9 residues (Fig. 3b). These included interactions with TM6 and TM12 (e.g. F336 and Q983, respectively) that mirrored paclitaxel, as well as contacts with TM1 (L65/L67/M68) and TM11 (M949/Y950/Y953) as well as TM5 (Y310), displaying wide contact profile on both sides of the cavity. Notably, In ATP/ADP (Fig. 3d), verapamil distributed the furthest toward the extracellular exit route, as the drug interacted with extracellular portion of TM6 (L332, F335, F336) while the contact residues also slightly shifted to TM5 side (Y307 34%). The other partially hydrolyzed condition (ADP/ATP) resulted in verapamil distributing to the opposite side of the cavity, as it remained stable near TM11 (F942/M949/Y950) (Fig. 3f). Interestingly, this ADP/ATP condition led to the most open configuration of the cavity, as verapamil R1 and R2 moieties were visibly less involved in contacting the cavity (Fig. S4.2b), while the number of cavity residues above 0.25 contact frequency was the lowest among all nucleotide conditions, at 13 (Fig. 1b). This indicates that the asymmetric nucleotide driven redistribution of drug position can lead to looser packing of bound ligand and facilitate opening of the cavity toward the extracellular space. Interestingly, the ADP/ADP state representing fully hydrolyzed condition led to verapamil forming stable contacts with a cluster of residues primarily on the center/NBD2 side of the cavity, involving TM5, 6, 7, 8 and 12. The fully collapsed cavity with verapamil present in the ADP/ADP condition suggests that OF-open conformation likely represents a state P-gp may adopt in pre-hydrolysis or partially hydrolyzed nucleotide conditions. Throughout the open-state simulations (Fig. S4.2b), the two aromatic rings of verapamil continued to dominate binding interactions with with F343 on TM6 or F983 on TM12. The dimethoxyphenyl groups formed transient hydrophobic contacts with nearby residues, but these did not show a strong dependence on nucleotide state.

Overall, we observe that the asymmetric nucleotide condition in OF-open P-gp is correlated with positional bias of the drug to locate toward the specific side within cavity, in the direction of TMD region on the same side as NBD that is bound to ADP representing a post-hydrolytic condition. This observation marks an interesting phenomenon that may explain the drug extrusion mechanism: As a nucleotide on one NBS is hydrolyzed after another, the respective cavity movement may shift the bound ligand in either direction of the cavity which likely destabilizes the largely unspecific hydrophobic network between the ligand and the cavity to favor the eventual egress of the ligand.

### 2.3 Ligand contact profile in closed conformation

The OF-closed conformation corresponds to an outward-facing P-gp state modeled on the ATP-bound E-to-Q mutant, thus catalytically inactivate human P-gp structure solved by cryo-EM [4]. The OF-closed conformation is characterized by a collapsed extracellular gate, the two halves of the transporter came together at the top, constricting the pocket. Docking in this human P-gp structure placed ligands toward the extracellular interface, in a shallow binding mode where especially for paclitaxel, most of the molecule was positioned outside the cavity (Fig. 4g/h). As expected, the overall drug–protein contacts in this conformation were more limited as both paclitaxel and verapamil interacted with fewer residues relative to the OF-occluded or OF-open. Importantly, differences between pre- and post-hydrolytic states were evident primarily in the behavior of paclitaxel, whereas interactions of verapamil changed less dramatically with nucleotide state in the closed cavity.

**Figure 4:**
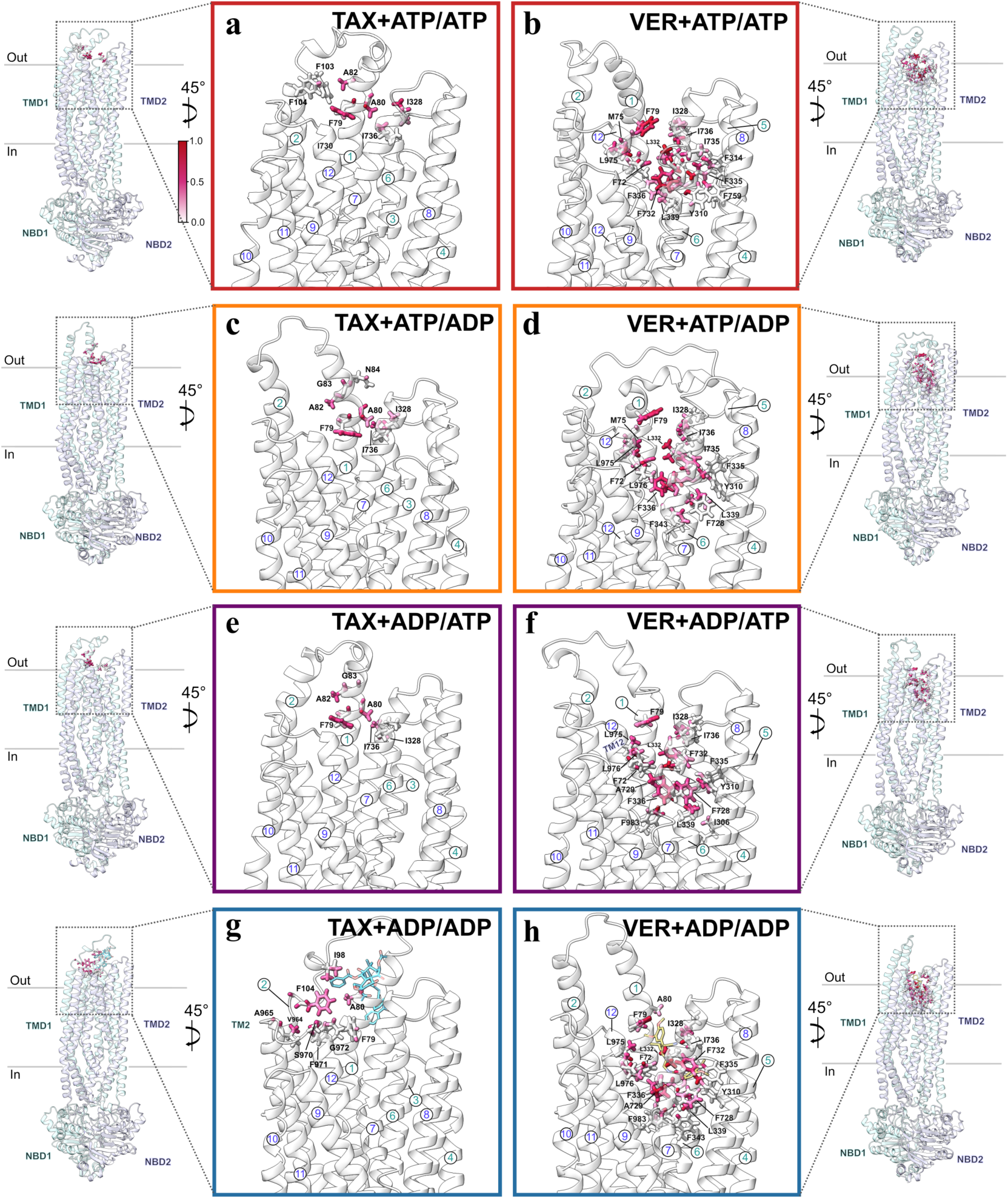
Residue level interaction frequencies between the TMD cavity residues and bound drug, either paclitaxel (TAX) or verapamil (VER) in OF-closed P-gp.(a-h) On the side, it shows a global orientation/representation of OF-closed P-gp to the side, with a box showing the clusters or TMD residues involved in the interaction with respective drug. The area shown with the box is displayed in an enlarged picture, where the residues with at least one atom above interaction frequency of 0.25 are shown with the side chain/backbone atoms colored in the respective frequency value. The initial docking position of TAX is shown in blue, in stick representation. Paclitaxel was docked outside the TMD ligand cavity, above TMD cavity near F79 side chain. (h) P-gp bound to VER with ADP/ADP. The initial docking position of VER is shown in yellow, in stick representation. Verapamil was docked further up in the TMD cavity than OF-open, which was near the interface between TMD cavity and extracellular solvent, near F336 and F72 side chains.

#### Paclitaxel in OF-closed conformation

Paclitaxel, as docked outside the cavity, had limited contact with the cavity as most of the interactions were with cavity gating residues F79, A80, I328 and I736 in all nucleotide conditions except ADP/ADP. Paclitaxel formed a stable contact with extracellular portions of TM1, 2, 6 and 7 as the drug remained close to the initial docked position. In contrast, ADP/ADP condition representing the fully hydrolyzed condition led to notable shift in the positioning of paclitaxel toward TM1/2/11 side of the extracellular side (still outside the cavity), as it formed low to moderate contact with the extracellular residues on TM1, 2, 11 and 12. Notably, some of the contacts seen in other nucleotide conditions were not observed (A80, I328, I736), suggesting the fully hydrolyzed nucleotide condition may further destabilize ligand contact at the edge of extracellular exit route and accommodate the drug near a temporary hydrophobic patch around TM1, 2 11, 12. Less consistent interaction with the cavity exit and establishment of interactions near TM1-2 loop away from the egress region in ADP/ADP condition could explain a potential mechanism of how P-gp prevents re-entry of drugs into cavity after both nucleotides are hydrolyzed.

#### Verapamil in OF-closed conformation

Verapamil, being smaller and more flexible, was able to be docked inside the small cavity available in the original cryo-EM structure [4]. Across the four nucleotide conditions, verapamil did not show any significant nucleotide-dependent shift in position within the cavity as verapamil interacted with a moderate set of residues (1̃5–16 with >0.25 frequency; Fig. 4b). The absence of positional shift in varied nucleotide suggests that the TMD cavity in OF-closed conformation is insensitive to nucleotides bound in NBS, thus likely represent a conformation that P-gp may adapt in post-hydrolysis condition.

### 2.4 Extracellular tunnel formation and TMD separation in OF P-gp

Using a 2.5 Å probe initiated from the central cavity (residues F336/F983), CAVER detected possible tunnels connecting the ligand-binding cavity of P-gp to external interfaces on the extracellular side and within the membrane. These tunnels were categorized by their exit region: channels 5a and 4a denote openings at the extracellular edges near the TM1/3 and TM7/9 interfaces, respectively, whereas 5b and 4b run toward the upper leaflet of the inner membrane near TM1/3 and TM7/9, and 6 leads into the extracellular space between TM6 and TM12. Notably, the OF-occluded conformation yielded no contiguous tunnel in any simulation, consistent with a sealed permeation pathway in the occluded arrangement.

In the OF-open conformation (Fig. 5), multiple egress tunnels from the drug-binding pocket to the periphery were identified, especially in the presence of paclitaxel. Paclitaxel-bound OF-open P-gp consistently exhibited a prominent tunnel along the TM7/9 interface (channel 4b) and a route toward the TM6/12 cleft (channel 6) in all nucleotide states, each remaining open in a majority of simulation frames (occupancies often 0.8–1.0 depending on nucleotide state). In addition, two tunnels on the opposite side of the cavity (near TM1/3) were intermittently accessible: an extracellular outlet between TM1 and TM3 (channel 5a) and a membrane-facing groove in the upper leaflet (channel 5b). By contrast, verapamil-bound OF-open simulations produced far fewer persistent tunnels. Under nucleotide conditions with ADP bound on NBS2 (ATP/ADP or ADP/ADP), no continuous tunnel was detected. Only in the NBS2-ATP bound states (ATP/ATP and ADP/ATP) did multiple pathways transiently appeared: in the ADP/ATP condition, all five tunnel types (4a, 4b, 5a, 5b, 6) opened at least sporadically, though with modest frequency (occupancies on the order of 10–40%). In ATP/ATP state, the TM7/9 and central cavity access was available in moderate occupancy (channel 4b and 6, 4̃0%) with rare accessible path on TM1/3 side (channel 5a 10%).

**Figure 5:**
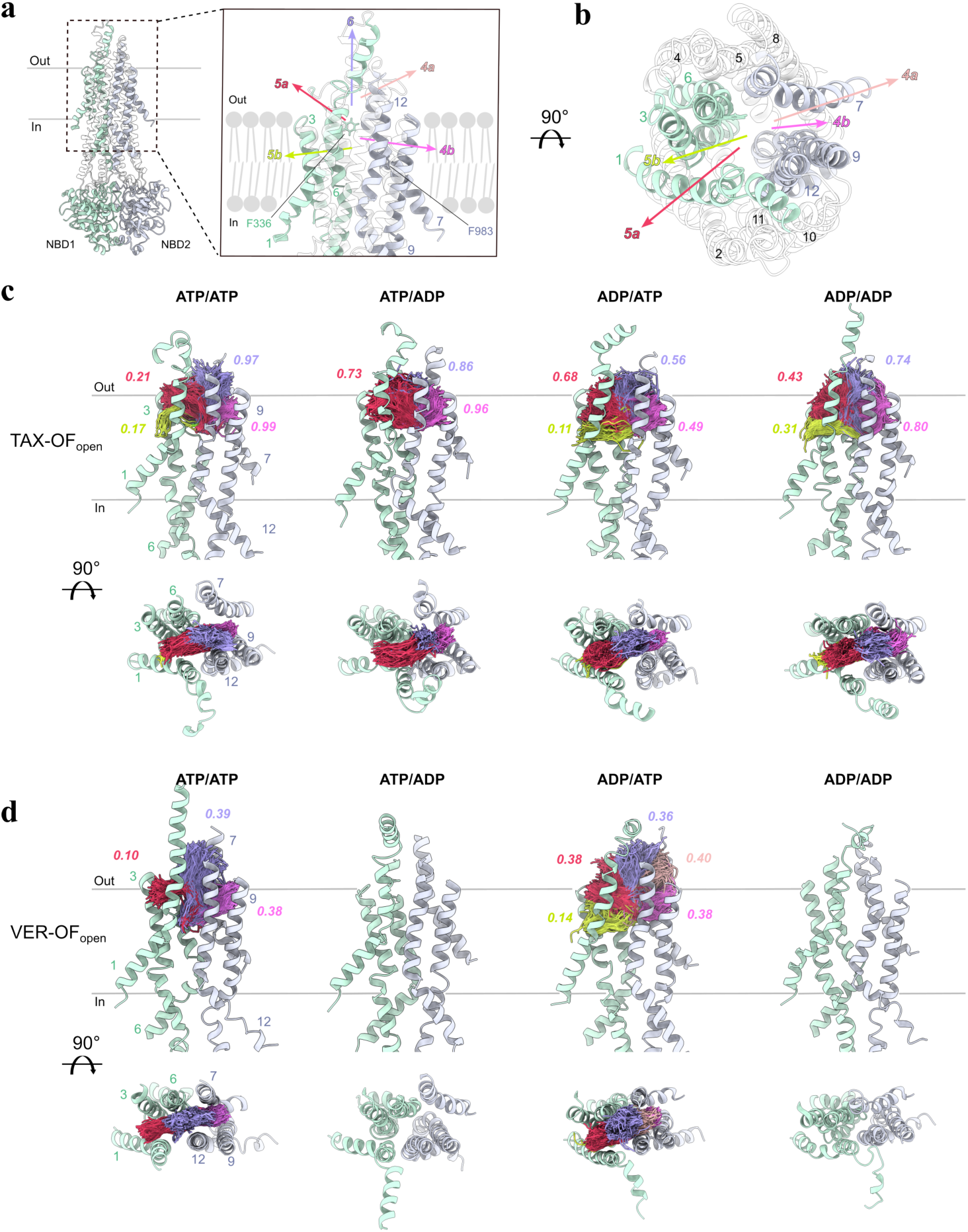
Potential ligand egress channels identified in OF-open P-gp bound with paclitaxel (TAX) or verapamil (VER) showing high structural plasticity upon nucleotide incorporation. (a) Viewed from the lipid bilayer cross section. Tunnels are defined as the following: 5a-extracellular side bordering upper leaflet access route via TM1/3 (green); 5b-upper leaflet access route via TM1/3 (green); 4a-extracellular side bordering upper leaflet access route via TM7/9 (blue); 4b-upper leaflet access route via TM7/9 (blue); 6-extracellular space egress toward the solvent near TM6/12. (b) Alternative view of the ligand access tunnels. (c) nucleotide-dependent ligand access gate formation in TAX bound OF-open P-gp configurations. Gray lines represent lipid bilayer boundaries to intra- and extracellular space. Relevant tunnel paths from all simulation replicas and corresponding tunnel occupancy are shown. TM1/3 (green) forms routes to tunnels 5a and 5b and TM7/9 (blue) routes to tunnel 4a and 4b. Last 70% of frames were used to analyze converged portions of the trajectories. Tunnels with occupancy values higher than 0.10 are depicted. (d) Tunnels detected in the VER bound OF-open P-gp configurations.

Thus, paclitaxel, a bulkier substrate, induces or requires a broader outward-facing opening that engages both the TM1/3-side and TM7/9-side exits, whereas the smaller verapamil tends to utilize a more limited egress route that is not consistently accessible except nucleotide conditions with NBS2-ATP bound state permits a transient opening. Overall, CAVER tunnels identified in OF-open suggest possible egress routes toward the lipid interface, especially for paclitaxel bound systems (only in limited nucleotide conditions for verapamil). Consistent accessible pathway between lipid interface and cavity center is likely to facilitate eventual diffusion of hydrophobic ligands like paclitaxel into the upper lipid leaflet/extracellular solvent interface.

The OF-closed conformation further underscored these ligand-specific differences (Fig. 6). With paclitaxel bound, no tunnels were identified in any nucleotide context, a finding consistent with paclitaxel remaining outside the cavity (Fig. 6a). This suggests that the ligand cavity in OF-closed conformation favors completely inaccessible cavity center from the extracellular space, ensuring the uni-directional transport. In contrast, verapamil-bound OF-closed simulations did permit an escape pathway in nucleotide states representing partial and fully hydrolyzed conditions (Fig. 6b). In both asymmetric nucleotide states with at least one ADP bound (ATP/ADP and ADP/ATP) and ADP/ADP, there was moderate occurrence of the central channel 6 and channel 4a toward extracellular solvent interface through the TM7/9 interface. Interestingly, there were no tunnels detected toward the lipid interface unlike the OF-open conformation, suggesting that OF-closed conformation even with the drug docked in the sequestered cavity does not open up toward the lipid milieu.

**Figure 6:**
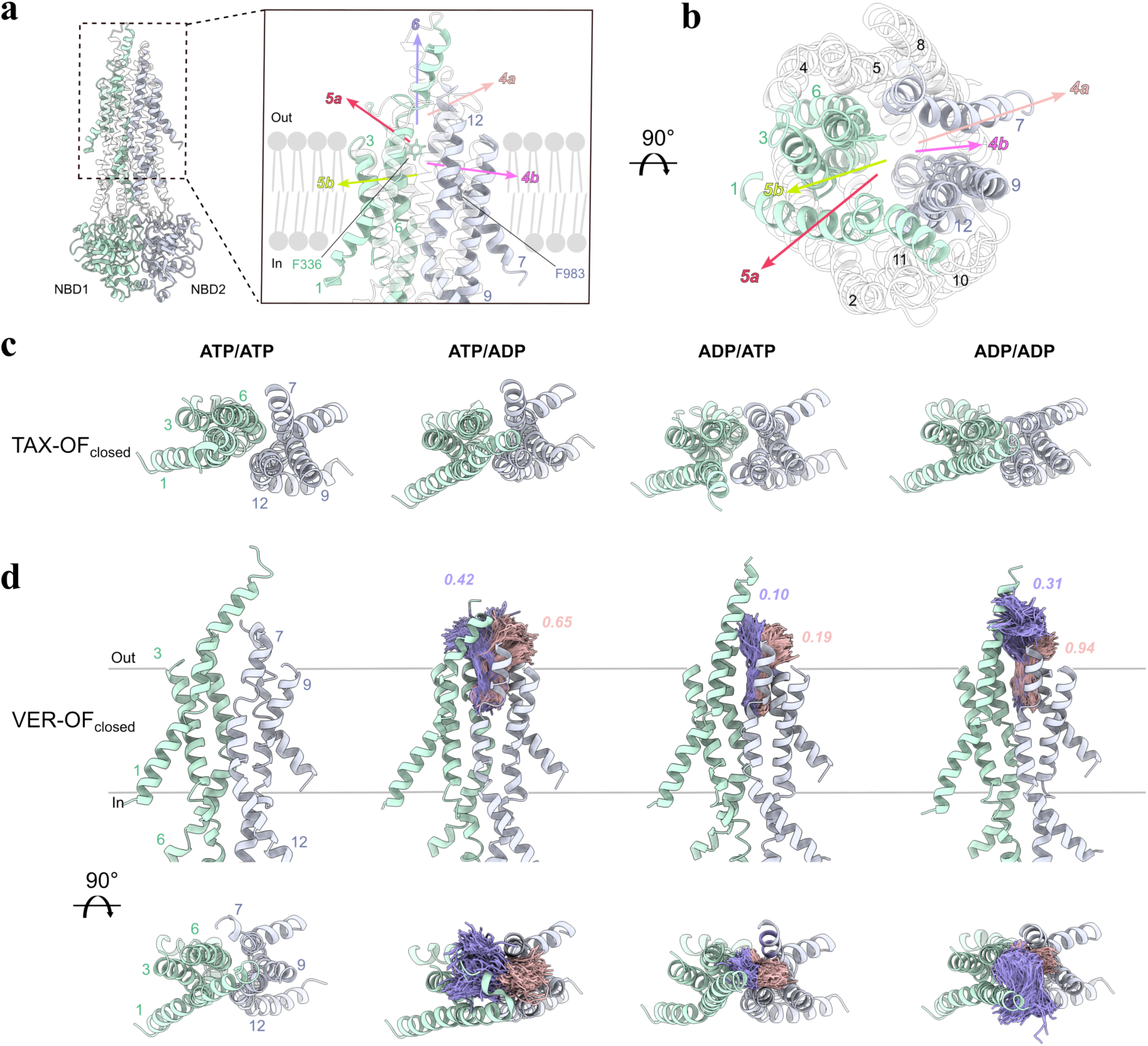
Potential ligand egress channels identified in OF-closed P-gp bound with paclitaxel (TAX) or verapamil (VER) showing high structural plasticity upon nucleotide incorporation. (a) Viewed from the lipid bilayer cross section. Tunnels are defined as the following: 5a-extracellular side bordering upper leaflet access route via TM1/3 (green); 5b-upper leaflet access route via TM1/3 (green); 4a-extracellular side bordering upper leaflet access route via TM7/9 (blue); 4b-upper leaflet access route via TM7/9 (blue); 6-extracellular space egress toward the solvent near TM6/12. (b) Alternative view of the ligand access tunnels. (c) Gray lines represent lipid bilayer boundaries to intra- and extracellular space. Relevant tunnel paths from all simulation replicas and corresponding tunnel occupancy are shown. TM1/3 (green) forms routes to tunnels 5a and 5b and TM7/9 (blue) routes to tunnel 4a and 4b. Last 70% of frames were used to analyze converged portions of the trajectories. Tunnels with occupancy values higher than 0.10 are depicted. (d) Tunnels detected in the VER bound OF-closed P-gp configurations.

### 2.5 DEER-referenced extracellular residue pair distance analysis

To quantify the conformational gating differences underlying these tunnel observations, we measured C*α*–Cα distances for five specific transmembrane residue pairs previously examined by Verhalen *et al.* using DEER spectroscopy (Fig. 7) [6]. These pairs, 329-972 (TM6–12), 320-955 (TM5–11), 209-852 (TM3–9), 209-955 (TM3–11), and 756-852 (TM8–9), span helices across the two halves of P-gp and report on the separation of key regions of the TMDs associated with the OF cavity shape and egress pathway formation. These extracellular pairs were selected because their separations in the outward-facing (OF) ensemble fall within the robust detection window of DEER technique ( 20–60 Å), enabling qualitative comparison to the experimental inter-spin distance distributions for P-gp [6]. As expected, the OF-open structures showed substantially larger separations for many of these helix pairs compared to the closed and occluded conformations (Fig. 7). For example, in paclitaxel-bound simulations the distance between TM5 and TM11 (residues 320–955) expanded to roughly 40 Å in the OF-open state, whereas it was constrained to about 35 Å in the paclitaxel OF-closed state and 30 Å in the occluded conformation. A similar pattern was observed for the TM6–12 pair (329–972) that delineates the extracellular cleft between the two halves: paclitaxel-bound OF-open models yielded a separation of 18–20 Å, roughly 1.5-fold larger than in the corresponding closed or occluded states (12–14 Å). Even the spacing between two adjacent helices within a single TMD bundle (TM8–9, residues 756–852) reflected the outward gate opening: with paclitaxel bound, the OF-open conformation showed TM8–9 distances around 32–33 Å, compared to only 25–27 Å in the OF-closed state and 24–26 Å in the occluded state. Each of these metrics confirms that the extracellular sides of the transporter diverge substantially in the outward-facing open conformation, particularly when a bulky ligand occupies the cavity.

**Figure 7:**
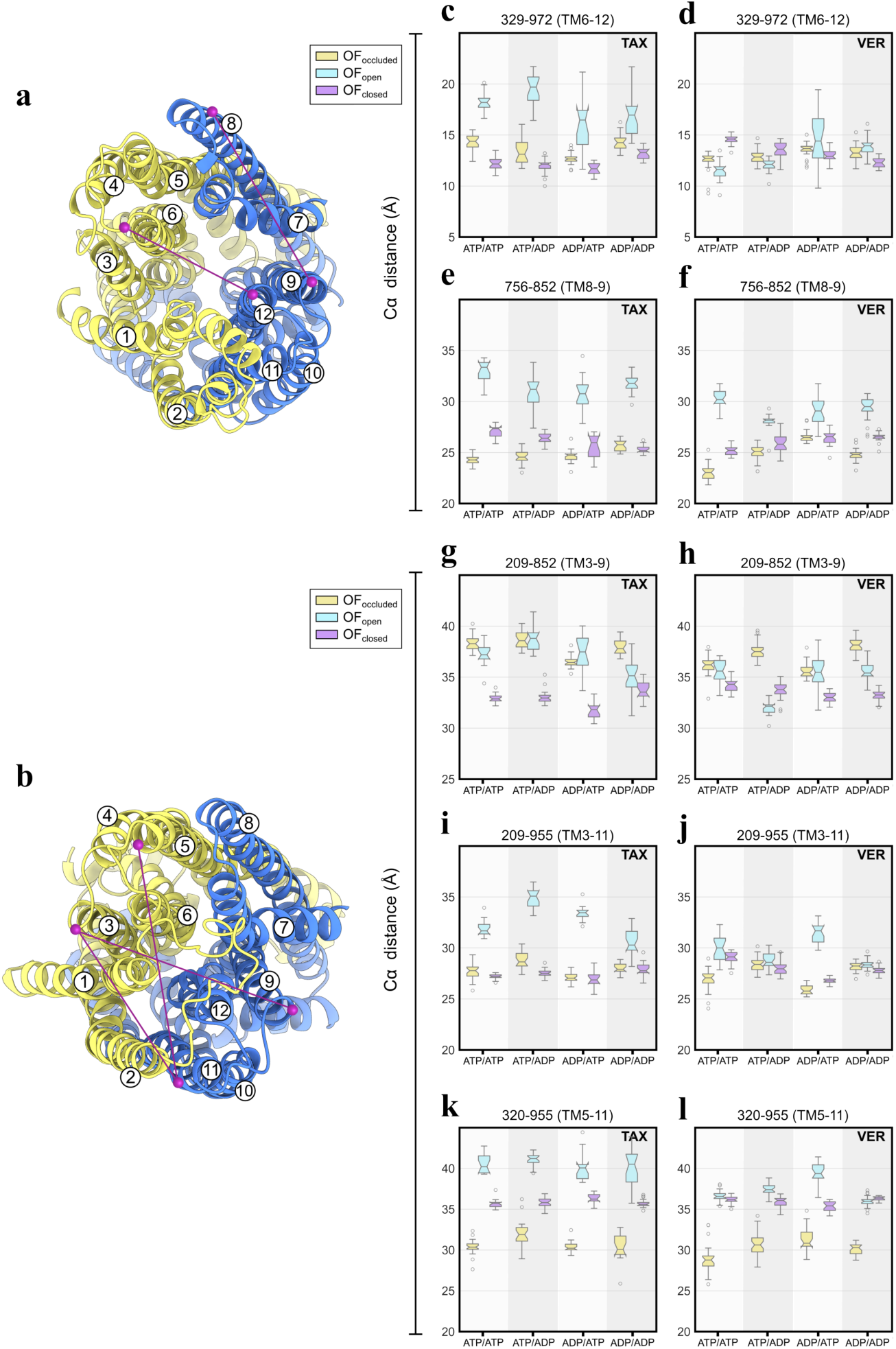
Box-and-whisker plots of residue pair distance distributions across nucleotide states in OF P-gp bound to paclitaxel (TAX) and verapamil (VER). (a,b) Top-down view of TMD, observing the OF P-gp down from the extracellular space. C*α* distances between residues. (c-d) For each panel, the x-axis is labeled with the nucleotide condition and the y-axis shows C*α* distances between the denoted residue pair. Within each nucleotide state, the distance distribution for OF-occluded, OF-open and OF-closed P-gp are colored in yellow, cyan and purple, respectively. Each box aggregates all frames from all replica trajectories for the given state, spanning the interquartile range from the first quartile (Q1) to the third quartile (Q3). The horizontal line inside each box is the sample median. The notch on each box visualizes an approximate 95% confidence interval for the median, computed as median ± 1.57 × IQR / n, where IQR = Q3 - Q1 and n is the number of observations contributing to that box. Whiskers extend to the most extreme data points within 1.5 × IQR below Q1 and above Q3. Observations beyond the whisker limits are shown as individual points (outliers).

Verapamil-bound simulations exhibited the similar qualitative trends in these distance metrics, albeit with generally smaller magnitudes. In the OF-open conformation with verapamil, the TM5–11 separation reached 37–39 Å under the ADP/ATP condition, modestly exceeding the 36 Å seen in the verapamil OF-closed state and well above the 29–30 Å measured in the occluded P-gp. By contrast, the effect of verapamil on the TM6–12 distance was less pronounced: the OF-open state yielded at most 14 Å between residues 329 and 972, similar to the 13–15 Å range observed in the verapamil-bound occluded and closed state. Notably, in the ATP/ATP-loaded closed simulation for verapamil, the TM6–12 spacing approached 14.6 Å, comparable to an open-state value, yet no tunnel was present under that condition, indicating that a partial outward splaying of these helices alone does not guarantee a ligand-accessible pathway. Meanwhile, the TM3–9 pair (209–852, spanning the opposite lateral side of the cavity) remained relatively wide in both open and occluded states (typically 35–38 Å) but narrowed to 33–34 Å in the closed states, suggesting that the overall spread between the two halves of the extracellular TMDs can remain further apart even in an outward-occluded, pre-ligand release conformation. In contrast, the diagonal TM3–11 distance (209–955) was above 30 Å only in NBS2-ATP bound nucleotide states. The TM3–11 distance above 30 Å correlates with the channel 5a occurrence from CAVER analysis which measures the pathway through TM1/3, thus it suggests that TM3–11 distance represents a relative open/closure dynamics of the cavity on TM1/3 side of the transporter. Taken together, these distance analyses corroborate the tunnel findings: conditions that produced open egress tunnels correspond to significantly enlarged separations of the relevant helices, whereas conformations lacking tunnels maintained tighter helix packing at the extracellular gate.

### 2.6 Interaction profiles between nucleotide moieties and NBS residues in drug-bound OF P-gp

We analyzed how a drug in the transmembrane cavity of the outward facing P-gp affected coordination of nucleotides at the two nucleotide binding sites (NBS1 and NBS2). The interaction profile of NBS residues are displayed in the heatmaps (Fig. 8), with residues labeled on x-axis and one of the three interaction types on y-axis. Key NBS residues from conserved motifs (Fig. 8a) were examined for interactions with ATP or ADP in four nucleotide states (ATP/ATP, ATP/ADP, ADP/ATP, ADP/ADP) across three OF conformations. The NBS residues involved include the Walker A lysines (K433 and K1076) that bind the nucleotide phosphates, conserved aromatic A-loop residues (Y401 and Y1044) that *π*-stack with the adenine base, the Signature motif “LSGGQ” loops (e.g. L531 from NBD1 and L1176 from NBD2) that contacts with the nucleotide ribose, the catalytic Walker B aspartates and glutamates (D555/E556 and D1200/E1201) and H-loop histidines (H587 and H1232) which interact near the phosphate groups, and charged residues from the intracellular coupling loops positioned at the NBD–TMD interface (Fig. 8a). The frequency of specific contacts, including salt bridges, P*_i_* stacking, and hydrophobic contacts, between each residue and the nucleotide was quantified (Fig. 8b,c).

**Figure 8:**
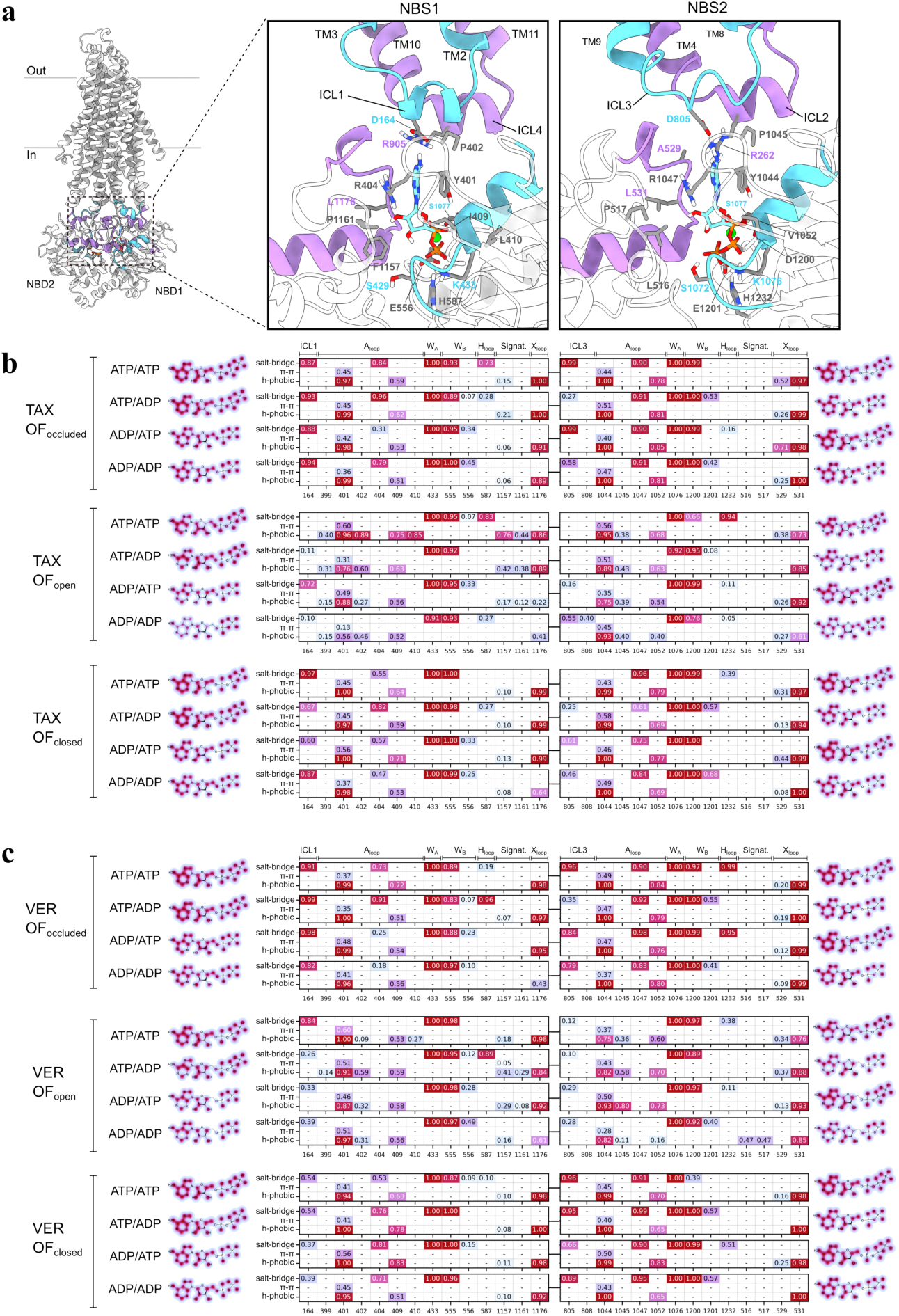
drug-bound at the extracellular portion of the TMD cavity in OF P-gp affects coordination between the NBS residues and the bound nucleotide. Interaction profile between the bound nucleotide and NBS residues were calculated. (a) The relevant NBS1 and NBS2 residues involved in the interaction are labeled, where the ICL1/3 and Walker A are colored in cyan and ICL2/4 and Signature motif are colored purple. (b) The interaction frequency of salt-bridge, *π*-stacking, and hydrophobic interactions between nucleotides and NBS residues in paclitaxel (TAX) bound OF P-gp, shown as heatmaps. The heatmap matrix is colored according to the interaction frequency for that residue and interaction type, normalized over all replica trajectories. Beside each heatmap, the respective nucleotide atoms involved in the interaction with NBS residues are colored in contour on top of the 2D chemical drawing based on the interaction frequency for that atom. (c) The interaction frequency heatmaps and 2D drawings, in the same format as explained in panel b, but for verapamil (VER) bound OF P-gp.

Overall, Walker A lysines K433 at NBS1 and K1076 at NBS2 showed near maximal salt bridge frequencies with nucleotide phosphates across all conditions. K433 and K1076 showed salt bridge frequencies at 1.00 in all nucleotide conditions, showing consistent coordination characteristic of OF P-gp conformation. Walker B aspartates (D555/D1200) and glutamates (E556/E1201) however, showed varied coordination frequencies, as the aspartates more consistently coordinated the nucleotide phosphate groups (9̃5% in most cases) while glutamates less frequently were involved in direct coordination of the phosphates. The catalytic H-loop histidine (H587/H1232) also showed dependence on NBD conformation for interactions with nucleotide phosphates as the few conditions with contact frequency above 0.5 occurred when coordinating ATP (with no exceptions).

A-loop residues that contact adenine showed measurable P*_i_* stacking with the base, and the frequencies varied with NBD conformation. Y401 at NBS1 and Y1044 at NBS2 showed P*_i_* stacking with adenine at approximately 0.40 to 0.50 in most conditions (Fig. 8). Few exceptions were found, including the TAX-OF-open P-gp in ADP/ADP condition, where *π*-stacking of adenine with Y401 was only 0.13 and the adenine and ribose sugar rings of NBS1-ADP were notably less frequently coordinated with NBS residues as seen in the 2D chemical drawing (Fig. 8a).

Residues from intracellular coupling helices showed frequent contacts with nucleotide moieties in both paclitaxel and verapamil bound P-gp. D164 (ICL1) at NBS1 and D805 (ICL3) at NBS2 showed high base associated contacts in most OF-occluded conditions, whereas OF-open P-gp showed inconsistent contact frequency for D164 and overall low frequency D805 across varied nucleotide states. In contrast, OF-closed P-gp showed a clear difference between the two drug-bound systems, as TAX-OF-closed systems showed overall moderate to high frequency nucleotide contact with D164 (60-97%) and low to moderate contact with D805 (0-61%), whereas VER-OF-closed systems had the opposite trend with low to moderate D164 contact (37-54%) and moderate to high D805 contact (66-96%). This drug dependent difference in ICL contact may reflect the absence and presence of a ligand within the cavity in paclitaxel and verapamil bound P-gp, respectively.

Elements contributed by the opposite NBD, including the Signature motif and the X-loop region, contributed to nucleotide binding in paclitaxel and verapamil bound P-gp. The X-loop residues L1176 and L531 forms a stable hydrophobic interaction with the bound nucleotide, often with high frequency (above 0.8), while the Signature motif residue contacts vary across the OF conformation and nucleotide states without a clear trend, showing the stochastic nature in nucleotide coordination even within the often thought static NBD sandwiched dimer in OF P-gps.

## 3 Discussion

The molecular dynamics simulation results provide new insight into how the outward-facing (OF) conformations of P-gp under various nucleotide states may orchestrate the late steps of substrate transport cycle. ABC exporters like P-gp undergo large conformational changes, yet fundamental questions remain about the timing of drug release relative to ATP hydrolysis and how unidirectional transport is favored. A key unknown factor is whether ATP hydrolysis occurs before, during, or after the substrate exits the transmembrane cavity. It is also unclear how the two nucleotide-binding sites (NBS) act, synchronously or in alternation, to power the conformational cycle [6]. Crystal and cryo-EM structures have captured P-gp and P-gp homologs in substrate-bound inward-facing and outward-facing states [5, 4]. However, fully outward-facing P-gp structures often lack bound drug due to low affinity within the cavity [5], and intermediate states are challenging to resolve due to high structural plasticity. DEER spectroscopy suggests that the outward-open state is transient and heterogeneous, appearing only upon or after ATP hydrolysis [6, 7]. Recent cryo-EM studies have trapped P-gp in occluded intermediates with substrate still in the cavity [9, 10], hinting at a multi-step release mechanism. Our simulations of drug-bound human P-gp in three OF conformations (closed, occluded, open) now tie these structural snapshots to nucleotide-dependent dynamics. Below, we discuss how nucleotide hydrolysis states influence the position of the drug in the cavity and potential egress pathways, how the NBS coordinate ATP in the presence of a substrate, and how these events may be coupled to yield an alternating, one-way transport mechanism. Unlike the previous Chapters, the flexible linker was left out of the discussion as it remains outside of the global domains of P-gp (due to its modeled location outside the NBD dimer interface).

### 3.1 Nucleotide state biases the position of the drug toward the extracellular exit

Across the OF-occluded, open, and closed systems, we found that introducing ADP (mimicking post-hydrolysis conditions) often caused the bound substrate to shift its contact profile in a way that suggests movement toward the extracellular exit. In the OF-occluded conformation, which has a sealed extracellular gate, paclitaxel and verapamil initially interacted deep in the central cavity (Fig. 2g,h). Notably, when both NBS contained ATP (the pre-hydrolytic ATP/ATP state), each drug formed relatively few high-probability contacts with the protein, indicating a looser binding pose (only 11 residues with >25% contact for paclitaxel and verapamil; Fig. 1a,b). By contrast, in nucleotide states containing ADP, especially the asymmetric ATP/ADP or ADP/ATP conditions, the drugs engaged a broader set of residues ( 15–17 contacts; Fig. 1a,b). For example, paclitaxel in the ATP/ADP state lost its unique ATP/ATP contacts in the lower cavity (Q195 on TM3, I299/A302 on TM5, F770 on TM8) and instead gained contacts nearer the extracellular side, such as F336 (TM6), F732 (TM7), and F978 (TM12) (Fig. 2a,c,g). Similarly, verapamil in the fully post-hydrolytic ADP/ADP condition made new contacts with Y310 (TM5) and F728 (TM7) high in the cavity (Fig. 2h), indicating the ligand had shifted slightly upward. These shifts suggest that once ATP is hydrolyzed in at least one NBS, the drug-binding pocket subtly rearranges to favor drug positioning closer to the extracellular vestibule. In other words, the hydrolysis transition correlates with the substrate moving along a “productive” pathway toward the exit.

In the OF-open conformation (with an intrinsically open extracellular vestibule), nucleotide asymmetry had a pronounced effect on ligand positioning. Paclitaxel and verapamil remained stably bound in the open cavity but redistributed laterally depending on which NBS carried ADP. Paclitaxel in a symmetric ATP/ATP state sat near the center-top of the cavity, contacting both halves of the protein (including TM6 and TM12, and TM1/TM7 at the portal; Fig. 3a). When ADP was introduced in one NBS, paclitaxel shifted toward the side of the cavity corresponding to the NBD that had ADP bound. Thus, in an ATP/ADP state (ADP in NBS2), paclitaxel drifted toward the TM5/TM7 side (NBD2 side of TMD) (Fig. 3c); conversely, in an ADP/ATP state (ADP in NBS1), it moved toward the TM11 side (NBD1 side of TMD) (Fig. 3e). Verapamil showed an even more dramatic nucleotide-dependent reorientation. In ATP/ATP, verapamil spanned the mid-cavity, interacting broadly with helices on both sides (TM6, TM12, TM1, TM11; Fig. 3b). With ADP in NBS2 (ATP/ADP), verapamil migrated toward the extracellular end of TM6 on the NBD2 side of TMD, engaging residues L332, F335, F336 near the outer opening (Fig. 3d). By contrast, with ADP in NBS1 (ADP/ATP), verapamil shifted toward the opposite side and settled closer to TM11/TM12 (NBD1 side), with fewer contacts overall (only 13 residues >25 frequency; Fig. 3f). This ADP/ATP scenario resulted in the loosest ligand packing (the two aromatic rings of verapamil partially lost contact with the protein), consistent with a substrate on the verge of dissociation. Fully post-hydrolytic conditions (ADP/ADP) stabilized verapamil in a more symmetric, tight cavity (Fig. 3h), resembling a collapsed cavity where the ligand was confined to the central pocket formed by TM5/6/7 and TM11/12. Taken together, these observations indicate that an asymmetry in NBS nucleotide, one ATP and one ADP, produces an asymmetry in the position of the drug. The substrate preferentially shifts toward the side of the cavity associated with the ADP-bound NBD, potentially reflecting the movements within the TMD that contributes to “squeeze out” the drug for eventual release. This finding echoes the “alternate access” pattern suggested by recent functional studies: P-gp may utilize two opposite egress routes in alternating fashion, each controlled by one half of the transporter [11]. In fact, rhodamine transport was shown to rely on two symmetric outer “gates” (Y310 on the N-terminal side; Y953 on the C-terminal side), either one of which can facilitate substrate exit [11]. Our simulations show nucleotide-dependent lateral movements of the drug toward the vicinity of Y310 or Y953, hinting at a similar drug expulsion mechanism. When one NBD hydrolyzes ATP, the drug is likely to be pushed closer to extracellular side as the preferable cavity contact shifts, presumably priming the less tightly bound substrate for release.

An interesting distinction emerged between the two drugs studied. Paclitaxel, being bulky and rigid, maintained strong contacts with a core set of hydrophobic residues (notably F343 on TM6 and Q986/F983 on TM12) in all conditions in all OF-occluded and OF-open conformers, and its multi-ring scaffold allowed it to simultaneously engage multiple helices (Fig. 2 and 4.3). This may have helped paclitaxel act as a “wedge” that partially props the cavity open, for example, paclitaxel-bound simulations frequently showed persistent openings of cavity toward the lipid interface and less collapse of the pocket even in ADP-bound states (Fig. 5c). In contrast, the smaller, flexible verapamil interacted with fewer residues at any given time and and the cavity tended to stay more closed (Fig. 5d). The binding footprint of verapamil narrowed significantly in the OF-open ADP/ATP condition (Fig. 3f), suggesting a less secure grip that could expedite its release. Experimentally, paclitaxel is a low affinity P-gp substrate that exhibits weak cooperative ATPase kinetics, whereas verapamil is transported more easily and stimulates ATPase in greater degree [16, 17]. Our data align with these behaviors: paclitaxel remained engaged with the cavity of P-gp across a range of nucleotide states, while verapamil became only transiently or loosely bound when the transporter adopted certain post-hydrolysis conformations. Indeed, the persistent contact of paclitaxel to the cavity (e.g., via TM6 and TM12 contacts) may necessitate a more drastic conformational change for full release, perhaps explaining why it is a relatively less robust substrate for P-gp with lower drug-stimulated ATPase activity [16, 18]. Verapamil, on the other hand, readily shifts and finds egress pathways with even subtle cavity openings, consistent with its efficient transport. Notably, verapamil has at least two known binding modes in P-gp, and occupancy of a secondary (low-affinity) site can inhibit transport [17]. The nucleotide-driven repositioning we observed might reflect verapamil transitioning between such modes: for instance, in the ADP/ATP state it may occupy only a primary site (hence fewer contacts), whereas in fully ATP-bound states it spans more interactions (as if engaging both sites or a broader interface). This highlights the polyspecific cavity of P-gp, which can accommodate substrates in multiple overlapping pockets [19]. Small chemical modifications are known to redirect compounds between these sub-pockets, drastically altering efflux rates [19]. Our simulations show that even without chemical changes, P-gp itself via nucleotide-dependent conformational changes can reposition a ligand between different pockets or orientations. This plasticity underscores how P-gp exploits both its own conformational dynamics and the flexibility of substrates to achieve transport.

### 3.2 Asymmetric ATP hydrolysis correlates with staged substrate release and NBD–TMD coupling

The interaction profiles at the NBS provide evidence that the two ATP sites of P-gp function in a coordinated but asymmetric manner. Throughout all simulations, the canonical nucleotide contacts were maintained: the Walker A lysine (K433 in NBD1, K1076 in NBD2) formed stable salt bridges to the, *β*- and *γ*-phosphates of ATP or ADP (Fig. 8b,c), and the Walker B aspartates (D555, D1200) consistently interacted with the nucleotide phosphates (often >90%). By contrast, the catalytic glutamate and histidine (E556/H587 in NBD1; E1201/H1232 in NBD2) showed nucleotide-dependent engagement. In ATP-bound sites, the H-loop histidine approached the *γ*-phosphate, forming a contact in a significant fraction of frames (50–60% when ATP was present, vs near 0% when that site contained ADP; Fig. 8b,c). This implies that the proper catalytic geometry, in which the histidine and the glutamate coordinate a water nucleophile and the *γ*-phosphate, was only attained in our simulations when an intact ATP occupied the site. Once ATP is replaced with one of the hydrolyzed product ADP, these interactions were lost, consistent with the idea that the post-hydrolytic NBS relaxes and the catalytic residues disengage from the nucleotide. This finding dovetails with quantum-classical simulations of BtuCD, which showed that the transition state for ATP hydrolysis requires a precisely positioned water, coordinated by the catalytic glutamate and a histidine, to attack the *γ*-phosphate, and that after phosphate cleavage the H-loop and glutamate rearrange [20]. The overall free energy change of ATP hydrolysis in the transporter was calculated to be near zero (G = +1.8 kcal/mol), meaning the chemical cleavage itself does not facilitate a major conformational change [20]. Instead, ATP binding is the key driver to tighten the NBD dimer and reorient the transmembrane domains, whereas hydrolysis serves to unlock the dimer for resetting [20]. Our data support this mechanism: in the presence of ATP (before hydrolysis), both NBS remain tightly engaged and the transporter stays in an OF state; once ADP replaces ATP, specific NBS contacts (e.g., H-loop/phosphate) break leading to less consistent coordination of ADP, which likely initiates the NBD separation and eventual return to inward-facing conformation.

Interestingly, we observed that when one NBS was in a post-hydrolysis (ADP-bound) state and the other still had ATP, the ATP-bound site often retained a catalytically competent configuration. For instance, in the ADP/ATP simulations, the NBD2 H-loop His1232 continued to interact with the *γ*-phosphate of ATP in NBS2, suggesting NBD2 remained ready to hydrolyze its ATP. Similarly, in ATP/ADP conditions, H587 of NBD1 stayed oriented toward the ATP in NBS1. This asymmetry suggests a sequential model: once one ATP is hydrolyzed (and its site loosens), the other NBS can still be poised to hydrolyze its ATP shortly thereafter. In effect, the two hydrolysis events may occur one after the other, not simultaneously, consistent with alternating catalysis models [6, 21]. Verhalen *et al.* deduced from spin-label distance distributions that, under turnover conditions, one NBS (likely NBS2) adopts an ATP-occluded conformation while the other is more open, implying only one ATP is tightly bound or hydrolyzed at a time [6]. Likewise, solid-state NMR of a bacterial ABC exporter trapped in a transition state analog showed one NBS with tightly bound post-hydrolysis mimic ADP·V*_i_* and the other NBS with a more weakly bound ADP [21]. Our findings mirror these findings: an ATP/ADP state structurally resembles a condition after one ATP has been hydrolyzed (ADP) and the second ATP is awaiting hydrolysis. Notably, in those asymmetric states we saw the substrate begin to disengage from the cavity (e.g., verapamil becoming loosely bound in ADP/ATP, Fig. 3f).

The simulations hint that drug–protein interactions could impart some bias on which NBS conducts ATP hydrolysis first. We observed subtle differences in how the two halves of P-gp engaged the nucleotide depending on the drug and conformation. For example, in paclitaxel-bound OF-closed simulations, the conserved aspartate in the intracellular coupling helix (ICL) of the side of NBD1 (D164 on ICL1) maintained stronger contacts with the nucleotide base than the equivalent residue on the side of NBD2 (D805 on ICL3), whereas with verapamil the reverse was true (Fig. 8b,c). This suggests that the presence of paclitaxel (which was largely outside the cavity in OF-closed) versus verapamil (still wedged inside the cavity) differentially affected in allosteric manner how each NBD engaged the nucleotide. Allosteric communication between the drug-binding pocket and NBS via the ICL coupling helices is well documented [13, 22]. Substrates or inhibitors bound in the TMD can alter the dynamics of the ICLs and NBD motifs, thereby favoring or disfavoring ATP hydrolysis. For instance, specific transport inhibitors have been shown to decouple NBD closure from TMD motions [22, 23]. In one study, the inhibitor zosuquidar locked NBD2 of P-gp in a more closed, protected state but caused atypical movements in the TMD and ICL4 [22], essentially uncoupling the normal concerted motion. By contrast, vinblastine binding stabilized certain NBD interactions in the post-hydrolysis state [22]. In our case, the two drugs we studied may likewise impose different allosteric constraints. The bulky presence of paclitaxel might symmetrize or stabilize the NBD dimer to some extent, whereas the smaller size of verapamil could allow more differential “tugging” on one side of the NBD–ICL interface. Consistent with this idea, verapamil in an ATP-bound state elicited asymmetric NBD arrangements in the cryo-EM structures [9] and is known to stimulate ATP hydrolysis in a dose dependent manner, suggesting it perturbs the two ATP sites unequally at high concentration [17, 18]. Ultimately, the simulation results support a model in which one ATP is hydrolyzed first (likely the one whose ICL connections are more stabilized by the current substrate conformation), and this partial action is enough to begin substrate release. The second ATP can then hydrolyze to drive the remaining conformational change and reset. This sequential hydrolysis ensures that at least one ATP remains bound at any time during the transport cycle to keep the NBD dimer (and thus the outward-facing orientation) intact until the substrate is expelled [6, 24]. Only once the drug is gone are both ATPs hydrolyzed, allowing the NBDs to separate and the protein to revert inward.

### 3.3 Outward gate opening is transient and coupled to nucleotide state, ensuring one-way flux

By analyzing ligand-accessible tunnels and key inter-helical distances, we found that opening of a continuous path from the cavity to the extracellular space occurs only under specific nucleotide conditions, and that these openings correspond to known outward-gating movements. In the OF-occluded conformation, as expected, no tunnel to the outside was detected in any simulation, regardless of nucleotide. This matches the structural definition of an occluded state: even with the NBDs dimerized, the outer helices (e.g., TM1 and TM11, TM6 and TM12) stay in contact, barring substrate exit. OF-occluded models likely represent an intermediate before final drug release, although the exact conformation of this intermediate state remains speculative as the initial model used to derive this conformation utilized covalent linking of ligand inside the cavity on mouse P-gp [10]. Indeed, our occluded simulations showed the substrate remained deeply embedded and fully solvent-shielded. Only when we simulated the OF-open conformation (based on Sav1866, which has a wide outer gap) did clear egress pathways appear. With paclitaxel bound, the OF-open P-gp maintained at least one open tunnel toward the extracellular side in virtually all conditions (Fig. 5c), unlike seen in ligand free OF-open simulations which showed nucleotide-dependent opening of the cavity (Fig. 2.4c). The most prominent path in paclitaxel bound OF-open P-gp was between TM7 and TM9 (channel 4b), leading into the upper leaflet of the membrane, and another between TM6 and TM12 (channel 6) opening directly to the extracellular aqueous phase. This suggests that a bulky substrate like paclitaxel inherently pries the TMDs apart enough to create egress routes independent of the nucleotide states. By contrast, verapamil-bound OF-open simulations showed nucleotide-dependent egress channel occurrence. In the pre-hydrolytic ATP/ATP state, some narrow openings appeared (e.g., transient channel 4b and 6 with 10–40% occupancy; Fig. 5d), and interestingly in the ADP/ATP state (ADP in NBS1) all tunnel types were intermittently observed (channels 4a/4b, 5a/5b, 6, albeit at modest frequency), similar to the observation in ligand free OF-open P-gp that showed the most channels all in high frequency in the ADP/ATP state (Fig. 2.4c). In verapamil bound OF-open, no continuous tunnel was found when NBS2 contained ADP (ATP/ADP or ADP/ADP), functionally occluded to the extracellular space and lipid interface. This is in contrast to the ligand free OF-open P-gp results, as ADP/ADP condition displayed minimal egress channel formations (Fig. 2.4). These findings imply that the outward gate of P-gp cavity can re-close even while the protein is in an overall OF conformation, especially if the substrate does not hold the gate open. In other words, the extracellular opening is conditional. For paclitaxel (large and pushing on the helices), the cavity access stays sufficiently open. However for the smaller verapamil, if the nucleotide state shifts to post-hydrolysis condition, particularly after one ATP is hydrolyzed on the NBS2, the cavity access to all sides tends to shut again, possibly trapping verapamil if it has not yet escaped. Functionally, this would correspond to a scenario where one ATP hydrolysis event is not enough to fully expel the substrate, leading P-gp to pause in an outward-occluded, drug still inside state until the second ATP is hydrolyzed. Such a state has been hypothesized and indirectly observed: P-gp likely enters a “doubly-occluded” intermediate after the first ATP hydrolysis, where the cytosolic side is closed (NBDs together) but the extracellular side has not fully opened [6]. Our verapamil ATP/ADP simulation fits this description, no tunnel, drug still bound, NBDs closed. Subsequent hydrolysis of the second ATP (simulated by ADP/ADP) then resulted in partial tunnel openings in the OF-closed system (discussed below), indicating the final power stroke to make the cavity more accessible.

The OF-closed conformation (modeled on ATP-bound E556Q mutant P-gp) offered a glimpse of the endpoint after substrate release. n paclitaxel-bound OF-closed simulations, no tunnels were detected in any nucleotide condition (Fig. 6a). This is consistent with paclitaxel being docked outside the cavity in this model due to the lack of cavity space in the initial structure. Interestingly, there was also no routes toward the lipid interface across all OF-closed systems unlike OF-open, where lipid-directed portals were common (Fig. 5c), indicating that even with a drug present the OF-closed cavity does not become accessible to the membrane. Mechanistically, this conformer could represent P-gp after expelling paclitaxel out of the cavity, where the extracellular gate “locks out” the substrate from re-entering. In verapamil-bound OF-closed simulations, where verapamil was initially placed just inside the cavity, we saw that under ATP/ATP the cavity was completely closed with no CAVER identified ligand pathway (Fig. 6c), consistent with a non-physiological, fully sequestered pocket imposed by the ATP-bound E556Q cryo-EM scaffold used for docking [4], which appears largely ligand-insensitive and likely represents a post-egress snapshot. However, in partially or fully hydrolyzed states (ATP/ADP, ADP/ATP, ADP/ADP) narrow tunnels appeared through the extracellular face (channel 6 between TM6 and TM12, or channel 4a between TM7 and TM9; Fig. 6d). These routes were not as persistent as those in the OF-open case, but their presence in ADP-containing states suggests that once at least one ATP have been hydrolyzed, the cavity access to extracellular solvent opens in varying degrees to potentially allow for the release of any remaining substrate. In other words, an OF-closed conformation with ATP bound (like the E556Q mutant structure) may still keep a small substrate sequestered if it had not diffused out during the initial opening. But upon hydrolysis (mimicked by ADP containing conditions in our simulations), the tight extracellular seal is breached, presumably the moment when the drug is actually expelled if it was still there. We suspect that OF-closed conformation is likely a post-ligand egress conformer due to our observation of the cavity closure in the absence of a ligand inside (paclitaxel bound P-gp and previously simulated ligand free OF-closed P-gp, see Fig. 2.4d), where hydrolysis on either side of the transporter may ensure the exit of a ligand if there is any present. This interpretation aligns with the idea that two ATP hydrolysis events are required for complete substrate egress: the first creates an outward-facing occluded state (drug accessible to exit but cavity not yet open, like in VER-OF-open in ATP/ADP state), and the second produces an outward-facing open state (like in VER-OF-open in ADP/ATP state with loose contacted drug with many cavity accessible channels observed, which is set for eventual drug egress), which then rapidly transitions to outward-closed once drug exits the cavity as P-gp resets back to the IF conformation [6, 25]. Likewise, single-molecule fluorescence and luminescence resonance energy transfer experiments on P-gp have suggested that the release of drug can coincide with the hydrolysis of ATP [26, 27, 28].

The distance measurements between specific transmembrane helices reinforce this model of transient, nucleotide-coupled gate opening. In the outward-open simulations, we observed significantly larger distances between helices that form the extracellular gate (Fig. 7c,d). Verapamil-bound systems showed a similar trend but to a lesser extent. Notably, in one case (verapamil, ATP/ATP) the TM6–12 distance reached 14.6 Å, as large as an open state, yet no tunnel was present, highlighting that a true permeation pathway requires more than just a single pair of helices separating. Other motions, like tilting or rotation of helices and unwinding of the extracellular loops, are needed to create a contiguous ligand egress channel [5, 10]. The DEER spectroscopy study of Verhalen *et al.* reported distributions for analogous distances in P-gp under different conditions, and those data showed that in the ATP hydrolysis transition state (trapped with ADP·V*_i_*) the extracellular gate distances were highly variable, some molecules had a wide opening, others remained closer to occluded [6]. This heterogeneity was interpreted as evidence of a short-lived outward-open state. Interestingly, our simulations capture a possible explanation: the ATP/ADP asymmetric state yielded some trajectories with a clear opening (verapamil in ADP/ATP had a wide TM5–TM11 and TM1–TM3 separation, for instance) while others remained closed, averaging to an intermediate distance. Correspondingly, we found that the TM3–TM11 distance (another reporter for the side portal between the TM1/3) was >30 Å only in states where NBS2 was ATP (Fig. 7d). This distance correlates with the occurrence of a tunnel on the TM1/3 side (channel 5a) in our CAVER analysis. In DEER experiments, the equivalent TM3–TM11 pair (human 209–955) showed a pronounced increase in distance in the presence of ATP plus vanadate [6, 7], consistent with one side of the outer gate opening significantly. Taken together, both the simulation and experimental distance measures support a scenario in which the extracellular gate opens asymmetrically and briefly as the transporter transitions through the post-hydrolysis state.

Preventing back-flow of substrates is crucial for an active transporter. Our results, in context with other studies, suggest P-gp has at least two mechanisms to enforce directionality at the outward facing state. First is a kinetic gate: the transporter does not dwell in a fully open state long enough for a substrate to diffuse back in from outside. The outward-open conformation exists only when ATP (or its analog) is bound and before hydrolysis resets the cycle, and even then it may flicker [6, 29]. Once hydrolysis occurs, the structure quickly either releases the drug or closes again. Second is a structural gate: physical “valves” formed by conserved residues that close in one orientation. For example, Kodan *et al.* identified in a homologous exporter a pair of conserved glutamines (in TM6) and paired tyrosines and lysines (in TM3/TM4) that form a seal in the outward-facing state, preventing substrate re-entry from the extracellular side [8]. Human P-gp contains analogous motifs (e.g., F72, F79 in TM1; F336 and F732 clear to the cavity-extracellular interface) which likely fulfill a similar role. In addition, our data suggests that TM1, a helix lining the outer cavity, acts as a flexible “lid”: it kinks open to allow egress and then straightens to seal the cavity once the substrate leaves. A recent cryo-EM of P-gp captured a substrate mid-exit and noted that the loop of TM1 formed transient contacts with the drug; upon release, TM1 returned to a straight conformation, occluding the route [10]. Our paclitaxel simulation is consistent with that model. Thus, the design of P-gp is to open just enough, just in time, to let the substrate out, and then immediately shut. This “spring-loaded” gate concept is supported by the behavior of the catalytically dead E556Q mutant: when it binds ATP, it reaches an outward-facing occluded state but cannot reset, and interestingly it shows an increased propensity to expose the UIC2 antibody epitope on the extracellular side [29]. That suggests the mutant samples an outward-open-like conformation, yet since it cannot hydrolyze ATP, it gets stuck within the OF conformation [29]. The wild-type P-gp avoids such a stall by using ATP hydrolysis as the trigger to close the outer gate and resetting the transporter.

### 3.4 Mechanistic model of P-gp transport: coupling sequential hydrolysis to alternating gate release

Integrating these findings, we propose the following mechanistic sequence for P-gp, which reconciles our simulation results with prior biochemical and structural data. In the inward-facing state, the large central cavity can bind substrates (often from the lipid bilayer) in multiple modes, as evidenced by our previous results discussed in Chapter 3 and by other experimental studies [19, 30]. Substrate binding increases the basal ATPase activity of P-gp, indicating allosteric crosstalk between TMD and NBD [25, 17]. When ATP binds to both NBSs, the NBDs dimerize, pulling the ICL coupling helices together and inducing a twist in the TMDs that collapses the inner cavity and begins to open the outer sides [8, 5]. Our OF-occluded simulations with ATP/ATP likely represent this pre-hydrolysis occluded state: NBDs tightly engaged, substrate now confined but not yet exposed to outside. ATP binding is thought to provide the free energy for this transition [20], essentially priming the transporter in an energized state with the substrate poised for export. At this stage, we saw that P-gp holds the substrate rather loosely (few contacts), suggesting the cavity has expanded or shifted in a way that decreases affinity (Fig. 1a,b). This is consistent with the observation that P-gp in an ATP-bound state has low affinity for many substrates [5]. The loose binding may be functionally advantageous, it means the substrate is ready to diffuse out as soon as a path opens. Notably, recent cryo-EM structures show that some substrates can bind at a “peripheral site” on the inner leaflet side of the cavity and then move to the central pocket during this transition [9]. That might explain why in our occluded ATP/ATP simulations the substrate had fewer contacts: it could be sampling a position closer to the gate, no longer snug in the inward-facing pocket.

ATP hydrolysis itself likely occurs when the substrate has destabilized one half of the NBD–TMD interface enough to allow slight water access and correct alignment of catalytic residues (H587, E556) at one of the NBS. In our simulations, conditions mimicking this single-hydrolysis intermediate (ATP/ADP or ADP/ATP) showed an intriguing dual effect: the protein remained outward-facing, but the substrate shifted markedly toward one side as the cavity became more accessible for potential egress (Fig. 5d). We interpret this as the moment of partial drug release. The side on which ATP was just hydrolyzed experiences a small conformational relaxation, the NBD dimer interface loosens asymmetrically [21, 6], which is transmitted to the TMD of that half. Consequently, the drug finds itself drawn to that opening half. Supporting this view, we saw paclitaxel and verapamil move toward the NBD2 side in ATP/ADP (ADP in NBS2) and toward the NBD1 side in ADP/ATP (ADP in NBS1), accompanied by loss of some drug–protein contacts (Fig. 3d,f). This likely corresponds to the substrate beginning to exit into the outer leaflet or extracellular vestibule. A recent time-resolved cryo-EM of P-gp with substrates did observe density for a drug moving toward the extracellular end in one half of the cavity, while still somewhat engaged in the other half [10]. Similarly, the two halves of P-gp can operate out of phase: one set of TM helices (TM1/3/11 side) might form an open gate while the other set (TM4/5/8 side) still clings to the substrate. Such a unilateral opening was directly visualized for a heterodimeric ABC exporter in an ATP analog-bound state [31]. Our asymmetric nucleotide simulations offer a mechanistic rationale: if only one NBD has hydrolyzed, only that side of the transporter can relax its grip. Interestingly, this state may correspond to the “twisted occluded” conformation described in some models, where the helices of one side are more outward than those of the other [24]. It is a vulnerable state because the substrate might slip out through the partially open cavity, which is desired, but it must not slip back in on the other side. The architecture of P-gp addresses this by maintaining the other ATP-bound NBS in a locked, closed state (preventing full collapse of the cavity) and by employing those one-way valve residues at the portal [8]. For example, if verapamil starts to exit via the cavity on the TMD region in the same side as NBD2, the opposite region of the cavity likely remains shut.

Finally, the second ATP is hydrolyzed, converting the transporter to a fully post-hydrolysis state (ADP/ADP + P*_i_*). This likely yields a brief access of the cavity in which the ligand can diffuse out. Any remaining substrate is expelled at this point. In our ADP/ADP simulations, we saw indications of this final release step: verapamil in OF-open ADP/ADP was found in a collapsed pocket with no continuous tunnels, suggesting the cavity has strong tendency to close up again, whereas paclitaxel, when docked just outside the cavity in the OF-closed ADP/ADP simulation, shifted to a very shallow, solvent-exposed position away from the cavity solvent exit (Fig. 4g). Within many replica trajectories of this condition, we observe paclitaxel fully diffusing away from the protein. The outward-open state in ADP/ADP state likely decays rapidly because both NBS now hold ADP and the NBD dimer can separate apart. That separation may be reflected in the TMDs, which eventually will reset the transporter back to the inward-facing state. Experimental support for this reset step comes from the catalytic E→Q mutant: when both ATP are bound but cannot be hydrolyzed, the protein is stuck outward-facing [4, 29]; but when hydrolysis proceeds normally, P-gp quickly returns to the inward facing state [32]. This is congruent with the paradigm that ATP binding drives the work cycle and ATP hydrolysis drives the recovery [20]. The inward-facing conformation is thereby restored, with ADP likely releasing (perhaps preferentially from one site before the other) to prepare for the next cycle [21, 33].

## 4 Conclusion

In summary, our simulations and analyses support a transport model in which P-gp harnesses ATP in a coordinated yet alternating fashion to extrude substrates. ATP binding brings the NBDs together and converts inward-open P-gp to an occluded state, squeezing the substrate. Sequential ATP hydrolysis then provides two small but critical nudges: the first hydrolysis partially opens the cavity to begin substrate egress, and the second hydrolysis completes the opening and then triggers closure of the outer gate behind the substrate. This “two-stroke” mechanism ensures a step-wise release of the more challenging, bulkier substrates and avoid back-transport, a principle of alternating access that is enforced in P-gp by the strict requirement of NBD dimerization (and ATP binding) for inner gate closure and by fast resetting after hydrolysis for outer gate closure. The interplay of drug movement and nucleotide state we observed lends credence to the idea that substrates actively tune the kinetics of the transporter. Some compounds, particularly bulky ones, may stall the cycle in the occluded state until both ATP are hydrolyzed, thereby coupling their own release to the full consumption of ATP, this is seen in the cooperative ATPase stimulation of paclitaxel. Other compounds might escape after none or only one ATP is hydrolyzed, potentially leading to partial uncoupling or less-than-maximal ATP hydrolysis, as can occur with certain substrates or at high drug concentrations that inhibit turnover. This model aligns with the “concerted but alternating” concept emerging from various studies, and frames P-gp as a finely tuned molecular machine that converts the chemical energy of ATP into a stepwise mechanical cycle to ensure substrates are moved out and kept out.

## 5 Materials and methods

### 5.1 Human P-glycoprotein 3D structural modeling

Cryo-EM structure of OF mouse P-gp bound to disulfide-linked AAC (PDB: 7zk6), OF Sav1866 crystal structure (PDB: 2HYD) [5] and cryo-EM structure of OF human P-gp (PDB: 6C0V) [4] were selected as templates for building full length human P-gp homology models. All sequences in template structures were mapped to human P-gp sequence (Uniprot ID: P08183) and missing residues were modeled using Modeller v9.23 [34]. A total of 5000 models of flexible linker (residues 630-699) were generated and a conformation with the highest Discrete Optimized Protein Energy (DOPE) was selected as starting configuration. The second starting configuration of the flexible linker was constructed by adopting the AlphaFold2 predicted structure (AF-P08183-F1-v4) [35].

### 5.2 MD simulation setup

Protein was embedded in 4:1 1-palmitoyl-2-oleoyl-sn-glycero-3-phosphocholine (POPC):cholesterol bilayer using CHARMM-GUI [36] based on the protein-membrane orientation predicted by OPM database [37]. Additional minimization was performed during system generation using CHARMM-GUI to relax the membrane lipids [36]. While a coarse-grained pre-equilibration route could further homogenize the bilayer, the present atomistic protocol yielded stable protein–membrane engagement across replicas and was sufficient for the purposes of this study. ATP-Mg^2+^ were docked by aligning nucleotide bound structure of ABCB10 (PDB: 4AYT) [38]. Simulations were performed using AMBER ff14SB force field for protein [39] and LIPID14 FF [40] for POPC-cholesterol membrane, and phosphate and magnesium parameters from Meagher *et al*. [41] and Allnér *et al*. [42]. The periodic TIP3P water box was used with Na^+^ and Cl^-^ ion concentrations of 150 mM. System was energy minimized using AMBER20 [43], applying harmonic restraints with a force constant of 1000 to 0 kcal/mol Å^2^ on the heavy atoms of protein, as described in other membrane simulation protocols [14]. The rest of the simulation protocol details are identical to the procedure described in section 2.6.2.

### 5.3 CAVER tunnel analysis

The CAVER 3.0 software [44] was used to identify the possible ligand access tunnels in the simulation trajectories with frames saved at intervals of 200 ps. Initial 30% of the frames of each of the trajectories were stripped to minimize starting conformation bias. All frames of the trajectories belonging to the unique P-gp conformer and nucleotide state were combined and aligned to an initial frame. For all CAVER analyses, tunnel calculation was performed excluding the residues 1-44, 370-710, 1013-1280 to only include the TMD region. The starting points were defined as the benzene carbons in F336 and F983 to represent the canonical ligand binding site. CAVER tunnel analysis was run using a probe radius of 2.5 Å, shell radius of 6.0 Å, and shell-depth of 4.0 Å.

