## Supplementary Figures for "Outward-facing P-glycoprotein bound to drug displays nucleotide-dependent drug egress mechanism"

### 6 Supplementary data

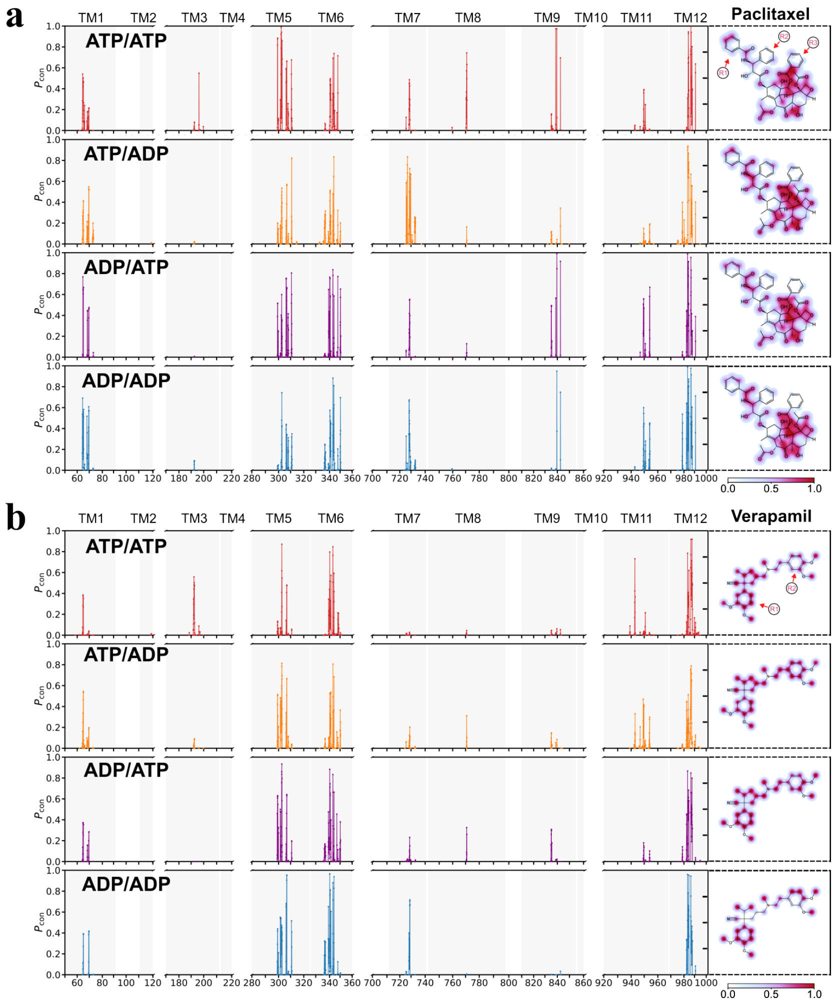


Supplementary Figure 1: Frequency of atom level interactions between the OF-occluded P-gp and the bound drug, either paclitaxel (TAX) or verapamil (VER) in the four nucleotide conditions, ATP/ATP, ATP/ADP, ADP/ATP, ADP/ADP, each colored in red, orange, purple, and blue, respectively. The ligand atoms involved in the interaction were colored with the respective interaction frequencies. (a) Frequencies calculated from the TAX bound OF-occluded P-gp. (b) Frequencies calculated from the VER bound OF-occluded P-gp.

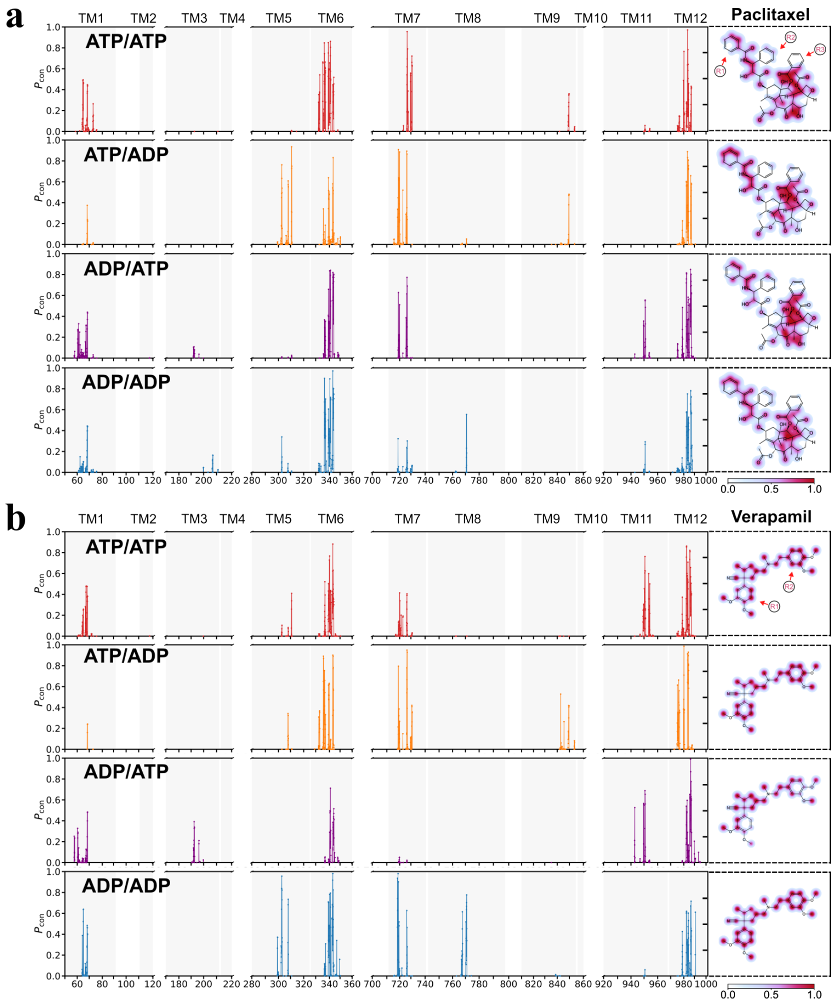


Supplementary Figure 2: Frequency of atom level interactions between the OF-open P-gp and the bound drug, either paclitaxel (TAX) or verapamil (VER) in the four nucleotide conditions, ATP/ATP, ATP/ADP, ADP/ATP, ADP/ADP, each colored in red, orange, purple, and blue, respectively. The ligand atoms involved in the interaction were colored with the respective interaction frequencies. (a) Frequencies calculated from the TAX bound OF-open P-gp. (b) Frequencies calculated from the VER bound OF-open P-gp.

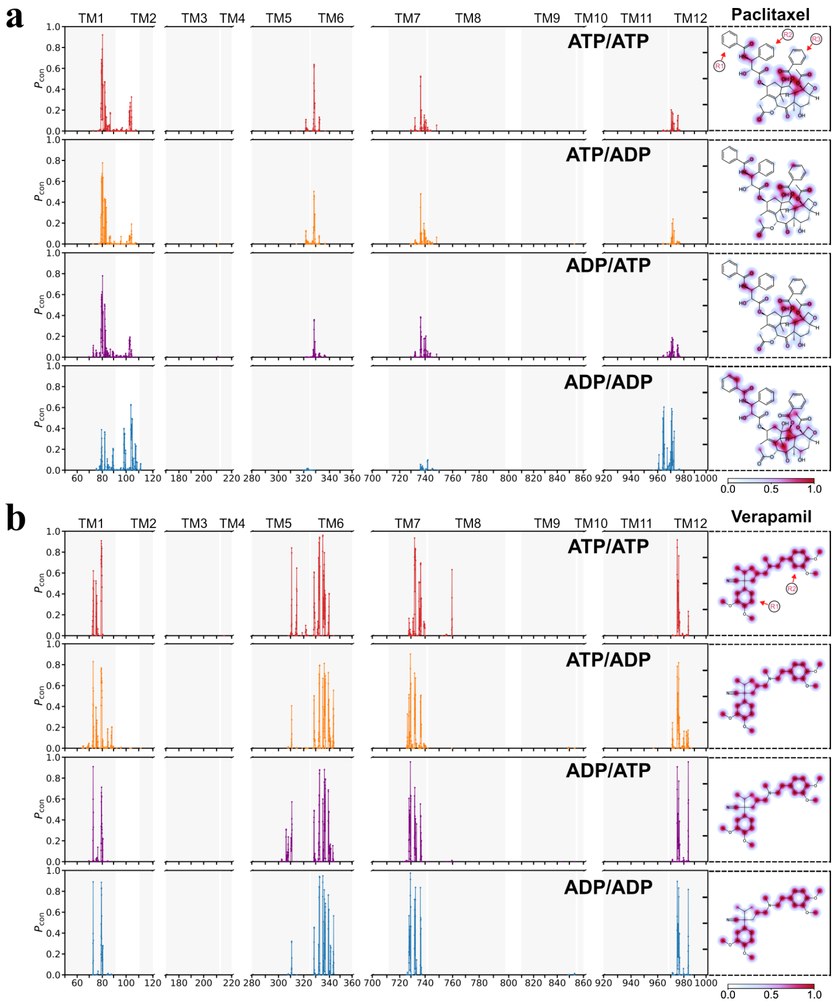


Supplementary Figure 3: Frequency of atom level interactions between the OF-closed P-gp and the bound drug, either paclitaxel (TAX) or verapamil (VER) in the four nucleotide conditions, ATP/ATP, ATP/ADP, ADP/ATP, ADP/ADP, each colored in red, orange, purple, and blue, respectively. The ligand atoms involved in the interaction were colored with the respective interaction frequencies. (a) Frequencies calculated from the TAX bound OF-closed P-gp. (b) Frequencies calculated from the VER bound OF-closed P-gp.
